# Mapping intratumoral myeloid-T cell interactomes at single-cell resolution reveals targets for overcoming checkpoint inhibitor resistance

**DOI:** 10.1101/2024.10.28.620093

**Authors:** Kate Bridges, Gabriela A. Pizzurro, Alev Baysoy, Janani P. Baskaran, Ziyan Xu, Varsha Mathew, Victoria Tripple, Michael LaPorte, Koonam Park, William Damsky, Harriet Kluger, Rong Fan, Susan M. Kaech, Marcus W. Bosenberg, Kathryn Miller-Jensen

## Abstract

Effective cancer immunotherapies restore anti-tumor immunity by rewiring cell-cell communication. Treatment-induced changes in communication can be inferred from single-cell RNA-sequencing (scRNA-seq) data, but current methods do not effectively manage heterogeneity within cell types. Here we developed a computational approach to efficiently analyze scRNA-seq-derived, single-cell-resolved cell-cell interactomes, which we applied to determine how agonistic CD40 (CD40ag) alters immune cell crosstalk alone, across tumor models, and in combination with immune checkpoint blockade (ICB). Our analyses suggested that CD40ag improves responses to ICB by targeting both immuno-stimulatory and immunosuppressive macrophage subsets communicating with T cells, and we experimentally validated a spatial basis for these subsets with immunofluorescence and spatial transcriptomics. Moreover, treatment with CD40ag and ICB established coordinated myeloid-T cell interaction hubs that are critical for reestablishing antitumor immunity. Our work advances the biological significance of hypotheses generated from scRNA-seq-derived cell-cell interactomes and supports the clinical translation of myeloid-targeted therapies for ICB-resistant tumors.

## INTRODUCTION

Solid tumors comprise heterogeneous neighborhoods of innate and adaptive immune cells, in addition to cancer and other supporting cells.^1,2^ The immune cell-cell communication networks established in these tumor neighborhoods contribute to the level of antitumor immune activity and responses to cancer immunotherapy.^3^ Many studies attempt to infer intratumoral cell-cell communication networks from single-cell RNA-sequencing (scRNA-seq) data.^4^ Putative cell-cell interaction maps are constructed by quantifying the expression of genes that encode for corresponding ligands, receptors, intermediate signaling proteins, and intracellular targets across interacting cell types. Despite these efforts, knowledge of how communication between immune cell subsets is rewired in tumors after immunotherapy remains limited.

Most cell-cell communication inference methods are limited in their quantification of potential ligand-receptor (L-R) interactions. Typically, L-R pairs are scored using averaged gene expression across cell types, or across pre-assigned clusters of single cells, and without spatial resolution.^4,5^ This discretization can complicate comparisons across conditions in which immune cells are in flux between cell states, e.g., following immunotherapy administration.^6,7^ Reliance on averaged gene expression also undermines the high granularity afforded by scRNA-seq and disregards the fact that, fundamentally, cell-cell communication occurs between individual cells. Understanding this communication at single-cell resolution is important to determine how therapy affects different immune cell subsets, specifically those which reside in distinct spatial neighborhoods within the tumor.^8,9^ Computational algorithms that infer cell-cell communication networks from scRNA-seq data at single-cell resolution have the potential to address this shortcoming,^10,11^ but such methods have not yet been tested as a means to discover changes in immune crosstalk induced by immunotherapy or combinatorial treatment schemes.

In this study, we developed a novel pipeline to analyze cell-cell communication networks derived from scRNA-seq at single-cell resolution to determine mechanisms underlying efficacy of combinatorial cancer immunotherapy with immune checkpoint blockade (ICB) and agonistic CD40 antibody (CD40ag). ICB that targets programmed cell death protein 1 (PD-1) and cytotoxic T-lymphocyte-associated protein 4 (CTLA-4) primarily acts on lymphocytes, inducing tumor cell-killing T cells.^12^ Despite demonstrated efficacy of checkpoint inhibitors anti-PD-1 (αPD-1) and anti-CTLA-4 (αCTLA-4) in many advanced malignancies, response rates range from 15-60% depending on tumor type, and many responders ultimately develop treatment resistance and subsequent disease progression.^13–16^ Drugs that target myeloid cells^17^ have been successful boosters for ICB because they promote interactions between T cells and myeloid cells, including dendritic cells (DCs) and tumor-associated macrophages (TAMs), that ultimately promote T cell anti-tumor activity and tumor regression.^18,19^ We and others have shown that treatment with CD40ag boosts T cell reinvigoration and improves anti-tumor responses to αPD-1, overcoming resistance in some cases.^20,21^ Further, CD40ag improved outcomes when combined with αPD-1 in patients with metastatic melanoma,^22^ and has been shown to contribute to tumor regression in a wide array of tumor models.^23–36^ Despite these recent studies, the CD40ag-induced changes in immune cell-cell communication that boost the antitumor response are incompletely understood, particularly as they concern the contributions of distinct, rapidly altered myeloid subsets to T cell stimulation and the ultimate reversal of ICB resistance.

Here, by analyzing single-cell-resolved, scRNA-seq-derived myeloid cell-T cell communication networks, we explored how myeloid-targeted immunotherapy drugs combine with ICB to rewire immune crosstalk and restore antitumor immunity in a preclinical model of melanoma. We demonstrated that changes in myeloid-T cell communication induced by CD40ag within the first 24 hours of treatment differ from those induced by ICB by differentially affecting distinct TAM subsets. We identified a suppressive Treg-macrophage IL-10 signaling axis associated with ICB resistance, and we show how ICB and CD40ag combine to break this suppressive interaction, while establishing durable interactions between DCs, macrophages, and T cells that are known to organize perivascular niches. We validate our findings with immunofluorescence and spatial transcriptomics. This work both advances the biological significance of scRNA-seq-derived hypotheses about cell-cell communication and supports the clinical translation of myeloid-targeted therapies for ICB-resistant malignancies.

## RESULTS

### Combining agonistic CD40 with immune checkpoint blockade yields durable overall survival in melanoma, allowing dose reduction of anti-PD-1 and anti-CTLA-4

To study how cell-cell interactions in the tumor microenvironment (TME) are altered with different immunotherapeutic drugs, we began by subcutaneously injecting WT C57BL/6 mice with YUMMER1.7 (YR1.7) cells, a melanoma cell line driven by BRAF^V600E^, CDKN2A^-/-^, PTEN^-/-^ mutations that was UV-irradiated to increase immunogenicity.^37,38^ We have previously shown that treating these tumors with CD40ag boosts their responses to αPD-1.^20^ Tumors were allowed to grow for seven days, and then cohorts of mice were treated with αPD-1 and αCTLA-4 at 200 μg/treatment/mouse (ICB hi) or 50 μg/treatment/mouse (ICB lo); CD40ag alone; or the combination of CD40ag + ICB lo. Treatments were repeated every 3 days for 5 total treatments, and mice were monitored for tumor growth and overall survival for comparison to untreated controls (Fig. 1A). While untreated YR1.7 tumors rapidly grow, most ICB hi-treated tumors regressed (Fig. 1B). In contrast, tumors treated with one quarter of the effective dose (ICB lo) largely continued to progress, with a median survival time of 21 days (Fig. 1B). CD40ag monotherapy achieved long-term survival in 38% of the mice and extended median survival time to 53 days. Importantly, CD40ag applied in combination with ICB lo resulted in tumor regression and overall survival in 71% of the mice (Fig. 1B). This is consistent with observations that CD40ag increases sensitivity to ICB in mouse models of pancreatic ductal adenocarcinoma.^26,30,33^ Of note, survival of mice treated with ICB lo and CD40ag was slightly superior to that of ICB hi-treated mice.

**Figure 1.**
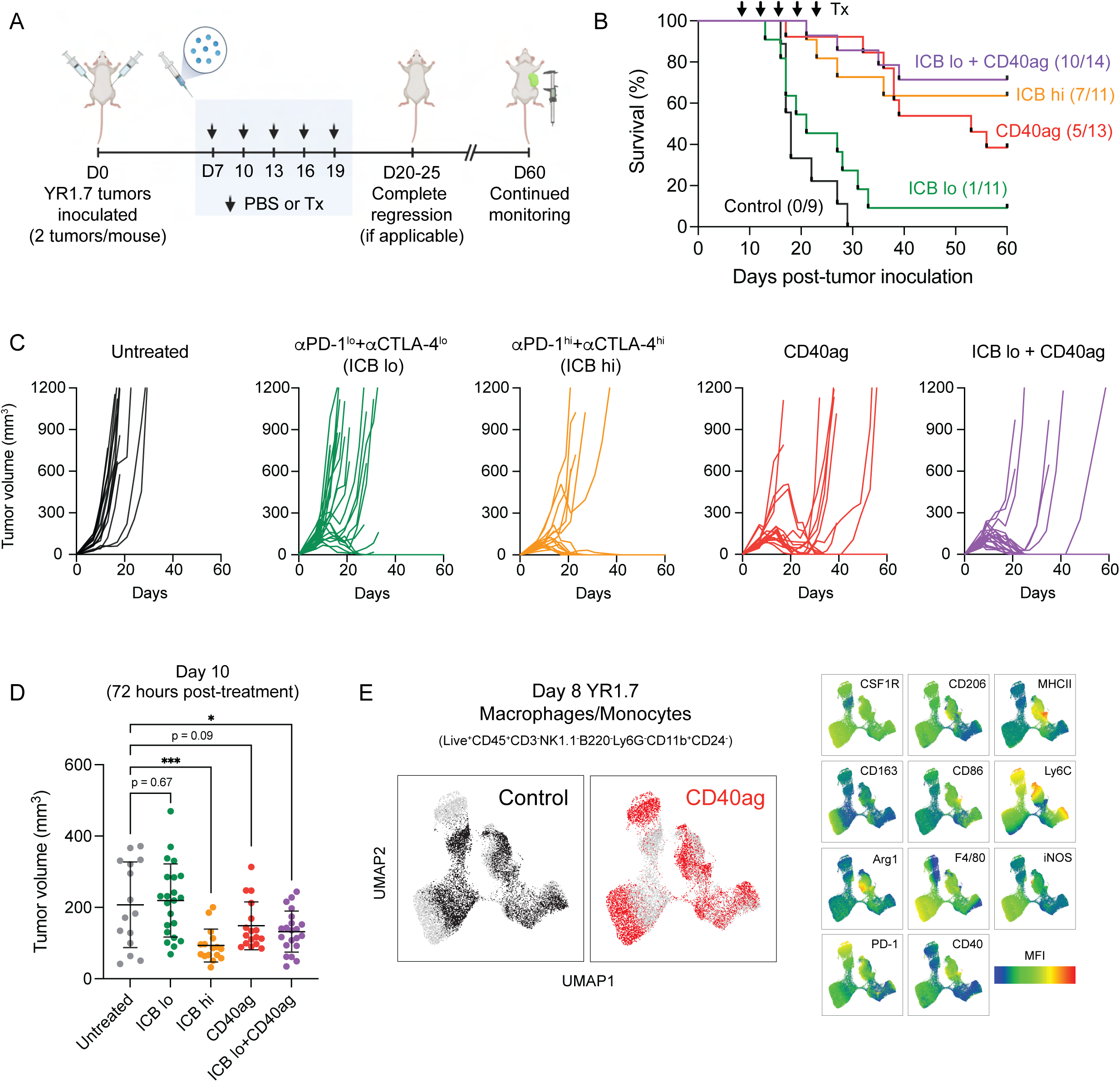
CD40ag combines with ICB to control growth of YR1.7 melanoma tumors. (A) Schematic describing treatment regimen for survival experiments. WT C57Bl/6J mice were inoculated s.c. with YR1.7 melanoma cells at day 0 (2 tumors per mouse). Treatments by were administered by i.p. injection and were initiated on day 7 and repeated every three days, for a total of five doses. Tumor growth was monitored through 60 days post-tumor inoculation. Created with BioRender.com. (B) Kaplan-Meier survival curves for YR1.7-bearing mice either left untreated (n=9, dark grey), or treated with high dose ICB (n=11, orange), low dose ICB (n=11, green), CD40ag monotherapy (n=13, red), or ICB lo + CD40ag (n=14, purple). (C) Corresponding tumor volume (mm^3^) curves over time for survival data shown in (B). (D) Tumor volume at day 10 (72 hours post-therapy initiation if applicable) for control, ICB lo-treated, ICB hi-treated, CD40ag-treated, and CD40ag+ICB lo-treated tumors. Summary data is represented as mean ± SD. *p < 0.05, ***p < 0.001 denotes statistical significance by one-way ANOVA with Holm-Sidak multiple comparisons correction (calculated in GraphPad). (E) Two-dimensional UMAP embedding summarizing expression of an 11-dimensional panel of macrophage phenotype markers measured by flow cytometry by macrophages and monocytes (Live^+^CD45^+^CD3^-^NK1.1^-^B220^-^Ly6G^-^CD11b^+^CD24^-^) sorted from 8-day YR1.7 tumors either left untreated (black) or treated with CD40ag (red) (data collection at 24 hours post-treatment if applicable). Plots to the left are colored by treatment condition, while the plots to the right are colored by mean fluorescence intensity (MFI) for each indicated surface marker, showing in detail the rapid phenotypic changes induced by CD40ag in TAMs.

CD40ag causes visible changes in YR1.7 tumors within 24 hours of treatment (Fig. S1A), consistent with our previous reports that CD40ag induces tumor necrosis and T cell mitosis within 48 hours of treatment, as well as an acute increase in circulating inflammatory cytokines and chemokines in both mice and human patients within 24 hours of treatment.^20,39^ Although CD40ag alone did not lead to complete tumor regression, it slowed tumor growth within 72 hours of therapy administration (Fig. 1C-D). In this same time frame, treatments with ICB hi or ICB lo + CD40ag induced significant decreases in tumor growth, while treatment with ICB lo alone did not (Fig. 1C-D). Moreover, we observed alterations in the tumor immune composition within 24 hours post-CD40ag monotherapy as assessed by flow cytometry (Fig. S1B-C), with increased representation of DCs and neutrophils. The monocyte/macrophage population was another predominant subset in the melanoma tumor immune infiltrate. Although its proportion did not change with treatment, we observed a CD40ag-induced phenotypic shift, with some monocyte/macrophage subsets exhibiting increased surface expression of functional markers including MHCII, CD86, iNOS, and Arg1 (Fig. 1E). Recent research has also reported an acute CD40ag-induced expansion of intratumoral neutrophils in models of colon and lung carcinomas.^25^ However, previous work by our group has suggested that CD40ag works independently of neutrophils in models of melanoma.^21^ Together, our data suggest that treatment with CD40ag rapidly rewires the myeloid compartment of the TME and overcomes ICB resistance to drive complete tumor regression and durable long-term survival. We were therefore interested in further exploring this acute stage of response to treatment.

### Single-cell RNA-sequencing (scRNA-seq) emphasizes heterogeneity in myeloid-specific responses to immunotherapy and highlights need for improved analytical strategy

Treatment with CD40ag acutely slowed tumor growth and ultimately improved survival outcomes in combination with ICB. To study rapidly-induced alterations in myeloid-T cell crosstalk which are associated with these emergent behaviors, we collected 8-day YR1.7 tumors for single-cell RNA-sequencing (scRNA-seq) 24 hours after administration of therapeutic regimens previously described (Fig. 1B-C). Prior to preparing cells for scRNA-seq, we enriched samples for T cells and myeloid cells (including DCs and macrophages) to better analyze acute treatment-induced changes in these populations (Fig. 2A). Additionally, we used barcoded hashtag antibodies^40^ to label individual tumors (Fig. S2A) and combined 4 tumors for each condition to increase the statistical power of our analyses.

**Figure 2.**
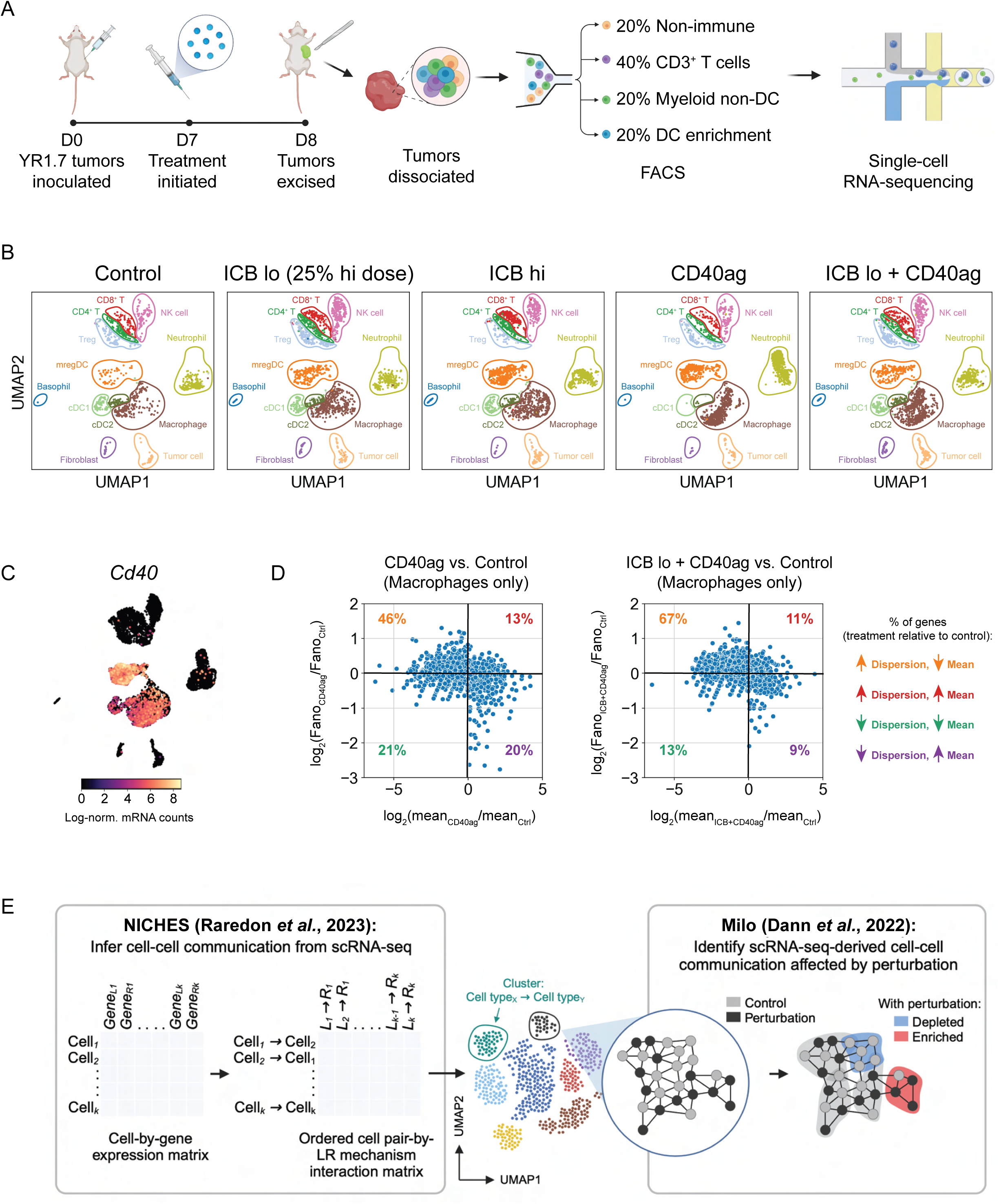
scRNA-seq emphasizes heterogeneity in myeloid-specific responses to immunotherapy. (A) Schematic describing treatment regimen and sorting strategy for single-cell RNA-sequencing (scRNA-seq) data collection. WT C57Bl/6J mice were inoculated with YR1.7 melanomas at day 0 (4 tumors from 2 mice per treatment group). Treatments were administered a single time at day 7, and tumors were excised and dissociated for sequencing 24 hours post-treatment at day 8. Prior to sequencing, tumors were FACS-sorted into four populations: CD45^-^ (tumor and stroma), CD45^+^CD3^+^ (T cells), CD45^+^CD3^-^CD19^-^XCR1^+^ and CD45^+^CD3^-^CD19^-^CCR7^+^ (DC subsets), and CD45^+^CD3^-^CD19^-^XCR1^-^CCR7^-^ (other myeloid cells). These four populations were then recombined at a 1:2:1:1 ratio and submitted for scRNA-seq. Created with BioRender.com. (B) UMAP embedding of control, ICB lo-treated, ICB hi-treated, CD40ag-treated, and ICB lo + CD40ag-treated scRNA-seq samples (n=4 tumors per group) separated by treatment condition and colored by cell type assigned by a neural network-based classifier. (C) UMAP plot as in (B) colored by *Cd40* expression. *Cd40* transcripts were primarily expressed by myeloid cell types, including macrophages and dendritic cell subsets. (D) Scatter plots comparing macrophage-specific log-fold changes in mean gene expression (x-axis) and Fano factor (y-axis) for CD40ag monotherapy (left) or ICB lo + CD40ag (right) relative to control. Each dot represents a gene that was highly variable (mean ≥ 0.0125, dispersion ≥ 0.5) in macrophages across treatment conditions (E) Schematic depicting our computational strategy for inference and comparison of scRNA-seq-derived cell-cell communication networks. We use the novel combination of NICHES for cell-cell communication inference and Milo for differential abundance testing to efficiently compare predicted interaction networks across experimental conditions and identify treatment-specific ligand-receptor interactions from scRNA-seq. Created with BioRender.com.

UMAP projection and supervised cell type annotation of the scRNA-seq data captured 12 cell types/subsets, including immune populations of macrophages, neutrophils, conventional type I and II DC subsets (cDC1s, cDC2s), DCs enriched in maturation and regulatory molecules (mregDCs), T cells (CD4^+^, CD8^+^, and regulatory T cells), and non-immune populations of fibroblasts and tumor cells (Fig. 2B). We observed visible transcriptional shifts in specific cell types across treatment conditions, most notably in the *Cd40*-expressing populations of macrophages and DC subsets (Fig. 2C). Interestingly, we further observed that macrophages from tumors treated with CD40ag-inclusive regimens occupied distinct regions of the larger macrophage cluster in two-dimensional space when compared to macrophages from tumors treated with ICB alone (Fig. 2B; in brown), revealing treatment-specific subsets.

While previous studies of CD40ag-induced changes in the TME have primarily implicated DC signaling to T cells as the major effector,^20,27,35^ few studies have discovered a role for macrophages. We hypothesized that this is in part due to the extreme macrophage-specific heterogeneity that we observe with CD40ag-inclusive treatment regimens. To quantify this CD40ag-induced, macrophage-specific heterogeneity from our scRNA-seq data, we calculated the mean mRNA counts and Fano Factor, or ‘noise-to-signal’ ratio, specific to each treatment condition for all genes that were variable (mean ≥ 0.0125, dispersion ≥ 0.5) across macrophages. In comparing these two metrics for both CD40ag-inclusive treatments relative to control, we found that noise increased with treatment for most genes (Fig. 2D), suggesting that individual TAMs respond differently to stimulation with CD40ag. Such differences are important in evaluating responses to immunotherapy and would be obscured by averaging across all TAMs.

To account for this heterogeneity in the myeloid cell response, we devised a novel strategy to generate and compare scRNA-seq-derived cell-cell communication networks without averaging across pre-assigned cell type clusters. We first use the Niche Interactions and Communication Heterogeneity in Extracellular Signaling (NICHES) algorithm^10^ to infer scRNA-seq-derived cell-cell communication networks at single-cell resolution (Fig. 2E, left). NICHES returns ordered cell-cell pair–by–L-R mechanism interaction matrices for each condition. To identify treatment-specific cell-cell communication axes, we then compare NICHES networks across treatment conditions using differential abundance testing with Milo^41^ (Fig. 2E, right). With this computational approach, we aimed to efficiently identify candidate extracellular interactions from scRNA-seq that underlie the responses observed with different combinations of immunotherapies, which might otherwise be overlooked by traditional cell-cell communication inference approaches.^4^

### Analyzing cell-cell communication at single-cell resolution uncovers candidate myeloid-T cell interactions that could limit efficacy of CD40ag monotherapy

We first applied our computational strategy to investigate cell-cell communication axes that were acutely modulated by CD40ag monotherapy. We focused these and all subsequent analyses on interactions between macrophages, DC subsets, and T cell subsets (Fig. 3A, top) because expression of genes encoding for therapy targets CD40, PD-1, and CTLA-4 and their cognate ligands were primarily limited to these cell types (Figs. 2C and S2B). NICHES networks for scRNA-seq samples from control and CD40ag-treated tumors included all nine possible directed cell type signaling pairs, as visualized in UMAP (Fig. 3A, bottom). These interactions could further be resolved into ‘sending’ and ‘receiving’ cell subsets, including T cell (CD8^+^, CD4^+^ non-Treg, and Tregs) and DC subsets (cDC1s, cDC2s, and mregDCs) (Fig. S3A, circled). For example, the larger macrophage-to-T cell cluster (Fig. 3A, bottom; in brown) visibly separated in two-dimensional space by the receiving T cell subset (Fig. S3A, right; in red, green, and blue), reemphasizing the need to infer L-R connections at the single-cell level to retain this information about heterogeneous signaling.

**Figure 3.**
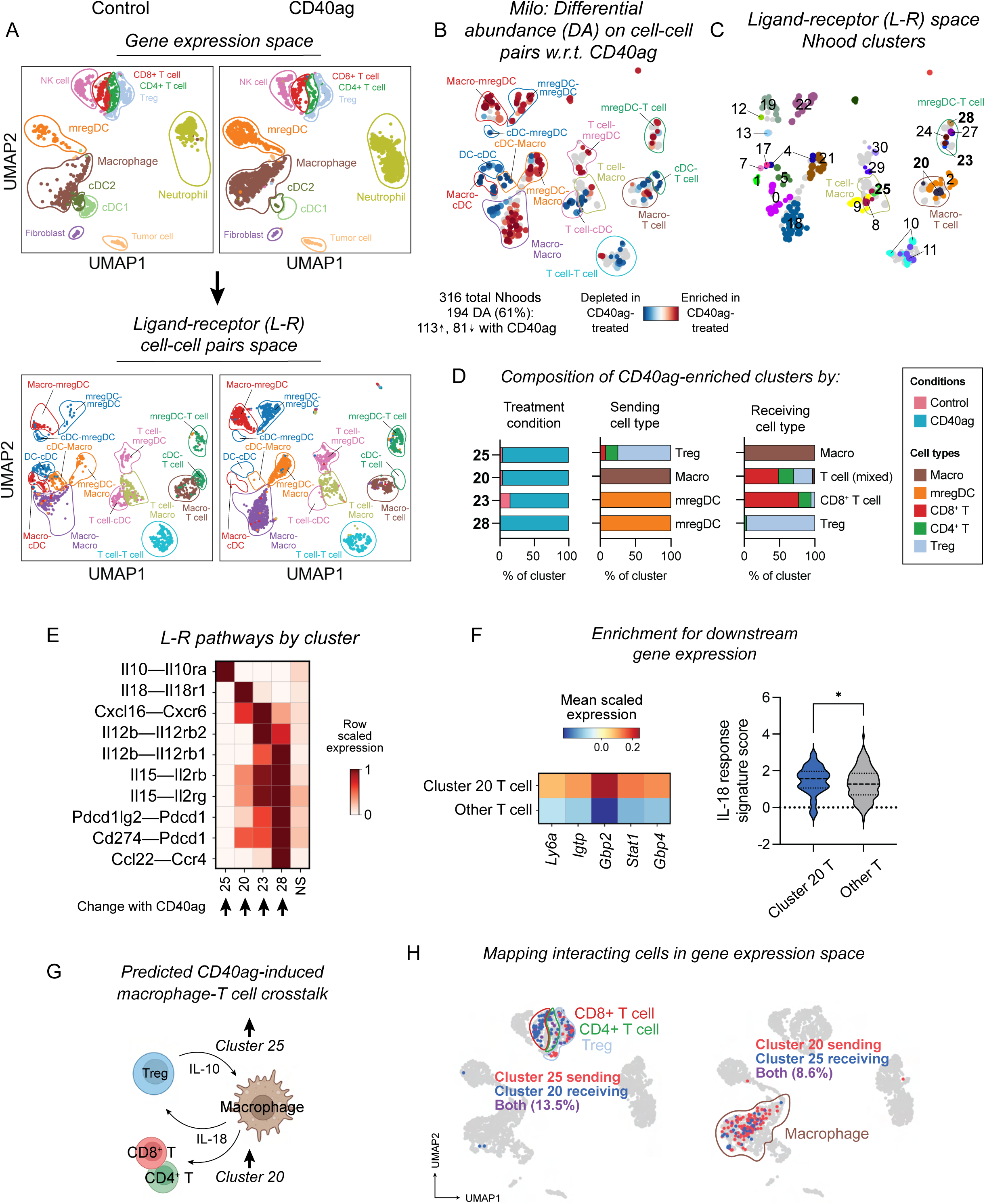
Analyzing cell-cell communication at single-cell resolution uncovers candidate interactions that could limit efficacy of CD40ag monotherapy. (A) UMAP embeddings of (top) single cells and (bottom) NICHES interactomes for control and CD40ag monotherapy-treated YR1.7 samples separated by treatment condition and colored by (top) cell type or (bottom) broad cell type pair. Cell type pairs are ordered to reflect cell type 1 (sender)-cell type 2 (receiver). (B) UMAP embedding of 316 Milo-assigned ‘neighborhoods’ (nhoods) of cell-cell pairs colored by differential abundance result w.r.t. treatment. Grey nhoods were not differentially abundant (n=127), while blue (n=81) and red (n=113) nhoods decreased and increased in frequency, respectively, with CD40ag treatment. (C) UMAP embedding of differentially abundant nhoods (n=194) colored by grouping for downstream analyses. (D) Stacked bar graphs summarizing the cell-cell pairs which make up differentially abundant clusters 25, 20, 23, and 28 by (left) treatment condition, (center) sending cell type, and (right) receiving cell type. (E) Heatmap of per-row scaled interaction scores for selected differentially predicted ligand-receptor axes (y-axis) across clusters 25, 20, 23, and 28 and compared to all non-differentially abundant neighborhoods (i.e., NS) (x-axis). (F) (Left) Heatmap summarizing mean scaled expression of the top 5 CD8^+^ T cell-specific IL-18 signature genes (via the Immune Dictionary). (Right) Violin plot summarizing distributions of single-cell-level average expression of the top 25 CD8^+^ T cell-specific IL-18 signature genes (via the Immune Dictionary) minus average expression of randomly sampled reference genes (i.e., ‘IL-18-response signature score’). Both plots compare expression or signature score for receiving T cells (cluster 20) versus all other T cells across treatments. *p < 0.05 denotes statistical significance by Mann-Whitney U test (calculated with GraphPad). (G) Schematic illustrating macrophage-T cell communication that we predict to be acutely upregulated with CD40ag monotherapy relative to control. (H) UMAP embedding as in (A, top) colored by T cells (left) and macrophage (right) predicted to interact in clusters 20 and 25.

To identify CD40ag-altered interactions after only 24 hours, we applied Milo to the 4,042 NICHES-derived cell-cell pairs to test them for differential abundance with respect to (w.r.t.) treatment. 194 Milo-assigned neighborhoods were identified as differentially abundant (Fig. 3B), which we grouped into 27 clusters for downstream analyses (Fig. 3C). Of these clusters, 14 were significantly enriched and 13 were significantly depleted with CD40ag treatment (Fig. S3B). A substantial number of clusters were between myeloid cell types (e.g., cDC1s to mregDCs), or between mixed cell types, which we did not examine in depth. We were most interested in how CD40ag established communication between macrophages and T cells or DCs and T cells, and therefore we focused on four clusters whose abundances significantly increased with CD40ag treatment (Fig. 3D): two involving mregDCs (28, 23) and two involving macrophages (20, 25).

mregDCs, also known as CCR7^+^ DCs, migratory DCs (migDCs), Lamp3^+^ DCs, or DC3s, have been described as both anti-tumor and pro-tumor.^42,43^ In line with this, Cluster 28 comprised mregDC (sender)-to-Treg (receiver) pairs, and cluster 23 comprised predominantly mregDC-to-CD8^+^ T cell pairs (Fig. 3D, center and right). These clusters were enriched for signaling via *Il12b*–*Il12rb(1,2), Il15–Il2r*(*b,g*)*, Cxcl16–Cxcr6*, and, specifically for cluster 28, *Cd274/Pdcd1lg2* (PD-L1/2)–*Pdcd1* (PD-1) and *Ccl22–Ccr4* (Fig. 3E). We and others have implicated DC-specific IL-12 in the mechanism for CD40ag in YR1.7 tumors^20^ and in other tumor models.^27,35,44^ IL-15 and CXCL16 have been shown to cooperate with IL-12 signaling in other contexts,^18^ but have not been noted as directly induced by CD40ag treatment. In contrast, the CCL22-CCR4 axis has recently been described to facilitate suppressive mregDC-Treg niches which sustain Treg activation and tumor progression.^42,45^ The observation that CD40ag affects both pro- and anti-tumor pathways suggests a possible limitation of monotherapy.

We also found that CD40ag treatment enriched for signaling between macrophages and T cells. Cluster 20 comprised macrophage-to-mixed T cell pairs, with enrichment for signaling via *Il18*–*Il18r1*, and to a lesser extent *Cxcl16*–*Cxcr6* and *Cd274*–*Pdcd1* (Fig. 3D-E). This cluster was of particular interest to us as we have previously uncovered a role for IL-18 signaling in YR1.7 tumor regression,^46^ although not in the context of CD40ag. To further support this putative interaction, we investigated whether there was evidence for activation of targets downstream of IL-18 receptor in receiving T cells. Using reference genes from the Immune Dictionary,^47^ we found that T cells predicted to receive IL-18 signal from macrophages in Cluster 20 were significantly enriched for expression of T cell-specific IL-18 signature genes (including, but not limited to, *Ly6a*, *Igtp*, *Gbp2*, *Stat1*, and *Gbp4*) compared to other T cells (Fig. 3F). Interestingly, we also predicted that CD40ag upregulates Treg signaling to macrophages via *Il10*–*Il10ra* (Figs. 3D-E; Cluster 25). IL-10-expressing Tregs have been previously shown to exert strong immunosuppressive effects on the cutaneous immune response.^48^

Our observation that CD40ag treatment increases both inflammatory and immunosuppressive signaling suggests potential mechanisms that could limit CD40ag efficacy (Fig. 3G). To explore if we would have been able to identify these axes using methods which rely on pre-clustered cell populations, we mapped the macrophages and T cells in Clusters 20 and 25 back into gene expression space (Fig. 3H). We found that the ‘sending’ and ‘receiving’ macrophages participating in these pairs overlapped, and so it is unlikely that they would have been separated into subsets prior to cell-cell communication inference. The T cells also overlapped, in addition to being drawn from distinct subsets (i.e., CD4 vs. CD8), emphasizing that key cell-cell communication axes could be mixed in terms of the sending and receiving cell type. This highlights the utility of looking for treatment-specific effects on highly resolved cell-cell communication pairs.

We further investigated if the cell-cell communication axes we predicted to be CD40ag-induced in YR1.7 tumors could be identified in another tumor model. We compared our CD40ag monotherapy-treated scRNA-seq samples to a publicly available scRNA-seq dataset which captured CD40ag-treated MC38 colorectal carcinomas 48 hours post-therapy.^25^ Similar to YR1.7, MC38 tumors exhibited a strong DC-specific increase in *Il12b* expression with CD40ag treatment (Fig. S3C-D). We applied NICHES to the dataset, but we could not perform differential abundance testing on the NICHES interactomes because the small number of replicate tumors per treatment condition underpowered the statistical comparison. However, directly probing the resulting interactomes for our connections of interest, we observed CD40ag-induced upregulation of the same myeloid-T cell signaling axes in MC38 tumors that we observed in the YR1.7 model. These axes included DC-to-T cell interactions along the *Il15–Il2r*(*b,g*) and *Cxcl16*–*Cxcr6* signaling axes, and macrophage-to-T cell *Il18*–*Il18r1* signaling (Figs. S3E-F). Importantly, an increase in the same immunosuppressive signaling axes were also predicted, including DC-to-Treg signaling through *Cd274/Pdcd1lg2* (PD-L1/2)–*Pdcd1* (PD-1) and *Ccl22–Ccr4*, and Treg-to-macrophage *Il10*–*Il10ra* signaling. Altogether, these analyses show that CD40ag treatment enhances both anti- and pro-tumor immune cell signaling between myeloid cells and T cells that is observed across tumor models. This provides strong evidence that our computational pipeline can uncover conserved cell-cell communication axes induced by immunotherapy.

### ICB induces acute, dose-dependent rewiring of intratumoral myeloid-T cell communication networks which may predict mechanisms of therapy resistance

We next wanted to predict which cell-cell interactions are modified by ICB within 24 hours of treatment and determine whether these were distinct from those induced by CD40ag. Further, because different doses of ICB yielded different tumor growth patterns and survival outcomes (Fig. 1B-D), we wondered whether possible mechanisms of therapy resistance would be observed at the lower ICB dose. To assess this, we used our computational strategy to analyze scRNA-seq samples from control, ICB lo- and ICB hi-treated tumors, collected as previously described (Fig. 2A-B), focusing on interactions between macrophages, DC subsets, and T cell subsets. Applying Milo to the resulting NICHES interactomes with dose as a continuous covariate (Fig. 4A), we identified 96 of the total 642 neighborhoods as significantly differentially abundant with respect to ICB dose, which we grouped into 33 clusters for downstream analyses (Fig. 4B-C).

**Figure 4.**
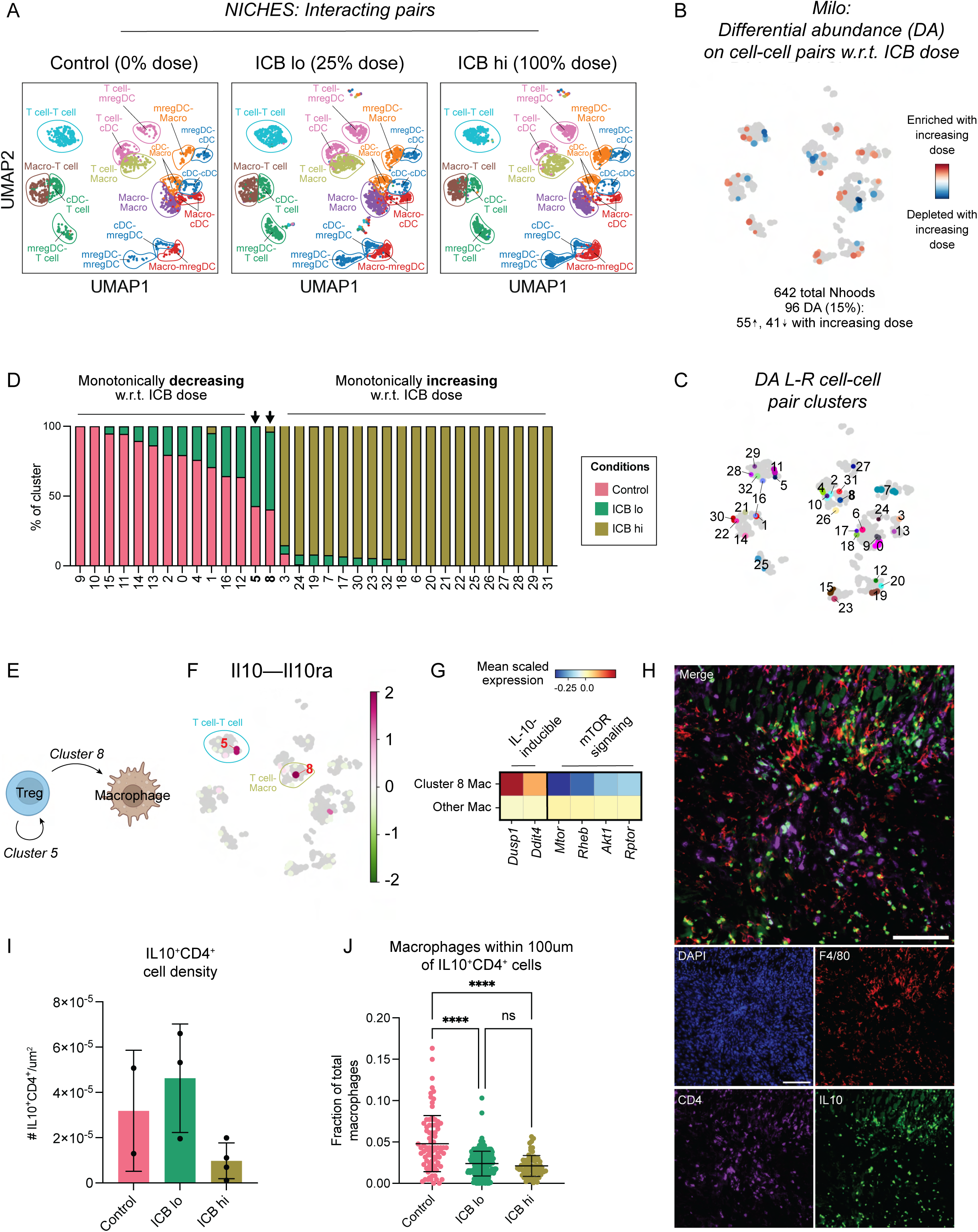
Immunosuppressive cell-cell communication between Tregs and macrophages in enriched with ineffective ICB dose. (A) UMAP embedding of NICHES interactomes for control, ICB lo- and ICB hi-treated samples separated by treatment condition and colored by broad cell type pair. (B) UMAP embedding of 642 Milo-assigned ‘neighborhoods’ (nhoods) of cell-cell pairs colored by differential abundance result w.r.t. ICB dose. Grey nhoods were not differentially abundant (n=546), while blue (n=41) and red (n=55) nhoods decreased and increased in frequency, respectively, with increased ICB dose. (C) UMAP embedding of differentially abundant nhoods (n=96) colored by grouping for downstream analyses. (D) Stacked bar graph summarizing the cell-cell pairs which make up each differentially abundant cluster in (C) by treatment condition. (E) Schematic describing Treg-to-Treg (Cluster 5) and Treg-to-macrophage (Cluster 8) communication predicted by our pipeline to be upregulated with ICB lo treatment relative to control and ICB hi. (F) UMAP embedding of NICHES interactomes grouped into Milo-derived nhoods and colored by predicted interaction score along Il10 – Il10ra as averaged across single cell-cell pairs in each nhood. (G) Heatmap summarizing mean scaled expression of IL-10-inducble genes *Dusp1* and *Ddit4* and mTOR pathway component-encoding genes *Mtor*, *Rheb*, *Akt1*, and *Rtpor* in receiving macrophages (cluster 8) versus all other macrophages across treatments. (H) Representative image of an 8-day untreated YR1.7 tumor inoculated in a B6.129S6-Il10^tm1Flv^/J mouse, stained for nuclei (DAPI, blue), F4/80 (red), CD4 (purple), and IL10 (GFP). Scale bars are 100μm. (I) Bar graph demonstrating quantification of CD4^+^IL-10^+^ T cell density (absolute number of CD4^+^IL-10-GFP^+^ cells divided by tumor area [μm^2^]) from control (pink), ICB lo- (green) and ICB hi-treated (yellow) tumor slices (n=2-4 regions from 1 tumor per treatment condition). Summary data is presented as mean ± SD. (J) Scatter plot demonstrating the fraction of total macrophages (F4/80^+^) within 100 μm of CD4^+^IL-10^+^ T cells from control (pink), ICB lo- (green) and ICB hi-treated (yellow) tumor slices. Summary data is presented as mean ± SD. ****p < 0.0001 by Kruskal-Wallis test (calculated in GraphPad).

These 33 differentially abundant clusters suggested that ICB rapidly changes immune crosstalk in the melanoma TME. 12 of these clusters were enriched in growing, untreated tumors and monotonically decreased in frequency as ICB dose increased (Fig. 4D). Many of these were enriched for immunosuppressive and anti-inflammatory-polarizing interactions. For example, Cluster 14 was enriched for macrophage-to-Treg communication via *Cd80*/*86*–*Ctla4*, while Cluster 0, comprising macrophage-to-macrophage interactions, was enriched for signaling through *Apoe* (encoding Apolipoprotein-E) to immunosuppressive receptors *Trem2*^49^ and *Lrp1*^50^ (Fig. S4A-B). In contrast, 19 clusters monotonically increased in frequency with ICB dose (Fig. 4D). These clusters were enriched for interactions known to induce T cell activation, trafficking, and effector function. For example, Cluster 25 was made up of mregDC-to-mixed T cell pairs predicted to communicate via *Cd80*/*86*–*Cd28,* and Cluster 30 comprised macrophage-to-CD8^+^ T cell pairs predicted to communicate via *Cxcl9/10*–*Cxcr3* (Fig. S4A-B). These findings are consistent with mechanisms of ICB efficacy,^51,52^ although few studies have demonstrated these changes at such early time points in the TME.

Notably, Cluster 5 and 8 were unique in that they were most abundant in ICB lo-treated tumors (Fig. 4D), suggesting that they might be enriched for interactions that are important for resistance to low-dose ICB. Cluster 8 primarily comprised Treg-to-macrophage pairs and Cluster 5 comprised Treg-to-Treg pairs (Figs. 4D-E and Fig. S4A), which were predicted to differentially interact via IL-10 (Fig. 4F), suggesting a potential mechanism of therapeutic resistance with ICB lo treatment. To build more support for this interaction, we investigated if macrophages in Cluster 8 expressed genes associated with activation of signaling pathways downstream of IL-10 receptor. Indeed, we found that, compared to other macrophages across treatments, cluster 8 macrophages had increased expression of IL-10-inducible genes *Dusp1*^53^ and *Ddit4*^54^ (Fig. 4G).

We further observed downregulation of mTOR pathway component-encoding genes in this cluster (Fig. 4G), which have been previously shown to be inhibited by IL-10-induced DDIT4 induction.^54^ Differential abundance on cell-cell communication pairs does not reveal if the change is driven by a change in the abundance of the cell type or in the cell signaling state. To consider this for Clusters 5 and 8, we explored how the abundance of cell subsets changed upon increasing ICB dose by applying Milo directly to the scRNA-seq data (Fig. S4C-D). This analysis revealed an enrichment for Tregs in ICB lo-treated samples (Fig. S4D-E; clusters 14 and 17), that expressed proliferative markers (*Mki67* and *Birc5*) and *Il10* (Fig. S4F). This finding is consistent with the predicted upregulation of IL-10 signaling in these tumors and suggests it is driven by an overall increase in Treg abundance with ICB lo. Interestingly, these Tregs also expressed higher levels of immune checkpoints including *Tigit, Tnfrsf18* (GITR), *Havcr2* (TIM3), and *Lgals1* (Fig. S4F), which can exert immunosuppressive effects on CD8^+^ T cells,^55^ and could be further contributing to therapy resistance and tumor outgrowth. Overall, our analysis predicts that while a high dose of ICB disrupts immunosuppressive signaling through IL-10, this signaling is maintained when a lower ICB dose is used, which might contribute to resistance.

To experimentally validate this prediction, we generated YR1.7 tumors in B6.129S6-Il10^tm1Flv^/J (IL-10-GFP reporter) mice^56^ to quantify the abundances and proximities of macrophages and IL-10-expressing Tregs in the YR1.7 TME and identify how these change with ICB treatment. At 7 days post-inoculation, we treated tumors with ICB at high or low doses or left them untreated. At 24 hours post-treatment, we collected these tumors and processed them for tissue imaging, with staining for GFP (*Il10*; green), F4/80 (macrophages; red), and CD4 (CD4^+^ T cells; purple) (Fig. 4H). While macrophage density remained largely constant across treatment conditions (Fig. S5A), we found that the density of CD4^+^IL-10^+^ T cells decreased with increasing ICB dose (Figs. 4I and S5B), in agreement with the abundances present in the scRNA-seq data. To estimate the extent of IL-10-mediated paracrine signaling between Tregs and macrophages, we measured the fraction of macrophages within 100μm radius of the CD4^+^IL-10^+^ T cells. ICB treatment, regardless of dose, significantly decreased the fraction of macrophages within 100μm of CD4^+^IL-10^+^ T cells compared to control (Fig. 4J). This trend held for a more stringent diffusion radius (Fig. S5C-D). Overall, in situ analysis supports our finding that achieving tumor regression with ICB requires a dose sufficient to decrease the number of CD4^+^IL-10^+^ T cells, which in turn modulates interactions with macrophages in the tumor.

### Combining CD40ag with low-dose ICB interrupts immunosuppressive cell-cell signaling while also establishing activating communication between myeloid cells and T cells

Thus far, our analyses have established that CD40ag and low-dose ICB individually rewire essential myeloid-T cell interactions in the TME in the first 24 hours after therapy, but each treatment on its own is not sufficient to overcome immunosuppression driven by signaling between myeloid cells and T cells. We were thus interested in understanding how the combined administration of CD40ag with low-dose ICB changes the landscape of cell-cell communication to improve survival outcomes relative to each treatment alone (Fig. 1B). Therefore, we again used our computational strategy to identify clusters of communicating cell-cell pairs that are enriched or depleted in ICB lo + CD40ag-treated tumors relative to control and ICB lo (Fig. 5A and Fig. S6A). From the NICHES networks, 60 Milo-assigned neighborhoods were found to be differentially abundant w.r.t. treatment efficacy (Fig. S6B), which we grouped into 20 clusters for downstream analyses (Fig. 5A, right; and Table S1). Of these clusters, 11 were enriched with combinatorial treatment (Fig. S6C).

**Figure 5.**
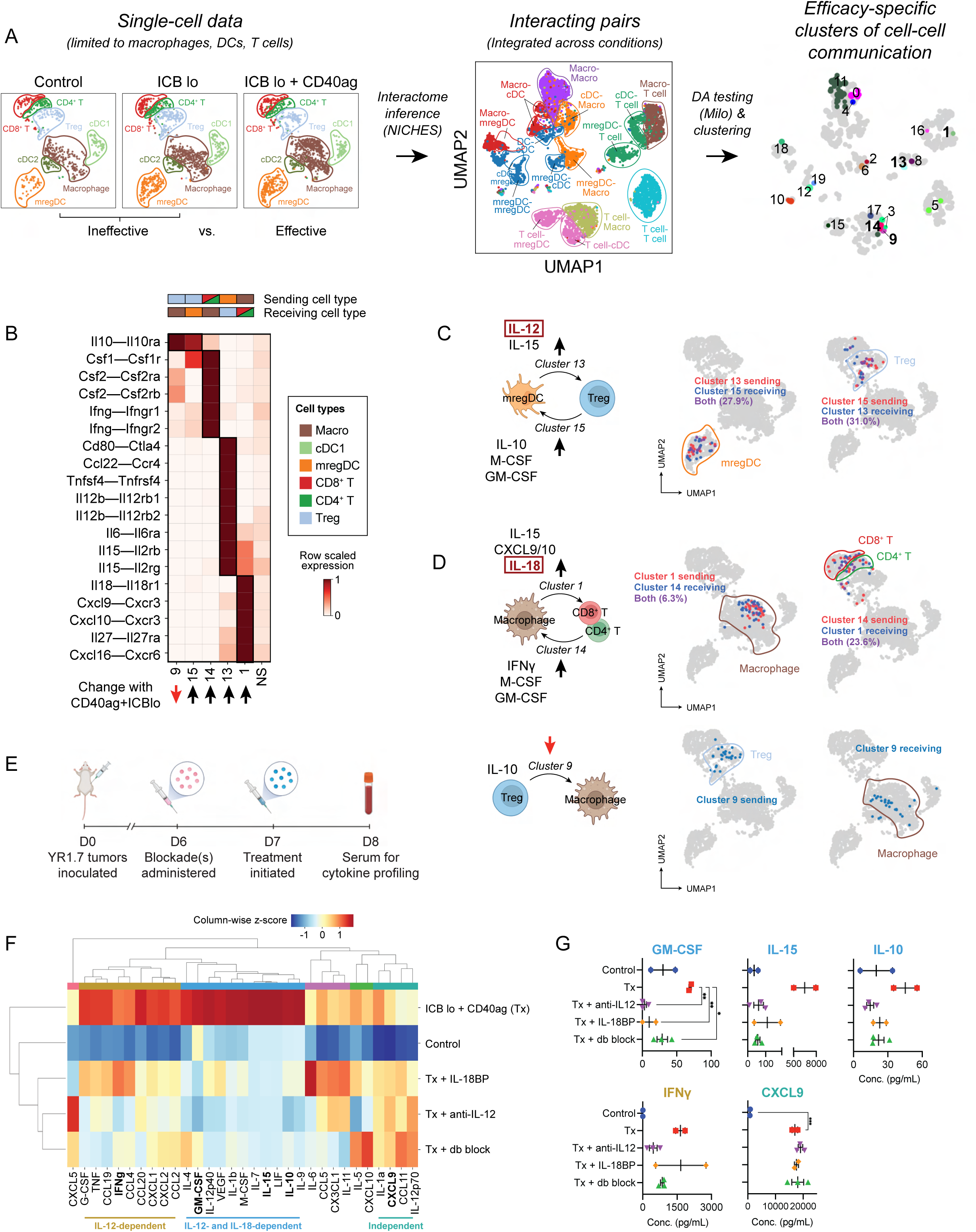
Single-cell resolved cell-cell interaction analysis shows that CD40ag combines with ICB to activate a coordinated cytokine-chemokine network between DCs, macrophages, and T cells. (A) UMAP embeddings of macrophages, DC subsets, and T cell subsets in gene expression space (left); and of corresponding NICHES interactomes (center) colored by broad cell type pair and (right) grouped into nhoods for differential abundance testing and colored by cluster for downstream analyses. Interactomes across experimental conditions (Control, ICB lo, ICB lo + CD40ag) are combined into a single visualization. (B) Heatmap depicting per-row scaled interaction scores for selected differentially predicted ligand-receptor axes (y-axis) across clusters 9, 15, 14, 13, 16, and 1 and compared to all non-differentially abundant neighborhoods (i.e., NS) (x-axis). The key above the heatmap distinguishes which cell type sender-receiver pairs are involved in each highlighted cluster. (C-D) UMAP embeddings as in (A, left) colored by (C) mregDCs (left) and Tregs (right) predicted to interact in clusters 13 and 15, (D, top) macrophages (left) and T cells (right) predicted to interact in clusters 1 and 14, and by (D, bottom) Tregs (left) and macrophages (right) predicted to interact in cluster 9. (E) Schematic illustrating the timeline of serum collection for peripheral blood cytokine and chemokine (C/C) profiling from ICB lo + CD40ag-treated YR1.7 tumor-bearing WT C57Bl/6J mice which had been previously treated with IL-12 and/or IL-18 blockade. Created with BioRender.com. (F) Hierarchical clustering of average C/C data from YR1.7 tumor-bearing mice either left untreated (n=3), or treated with ICB lo + CD40ag (n=2, abbreviated Tx in figure), Tx + IL18BP-Fc (n=2), Tx + anti-IL12 Ab (n=3), or Tx + anti-IL12 Ab + IL18BP-Fc (n=3). C/Cs were grouped into 6 modules for downstream analyses. (G) Example plots of peripheral blood cytokine expression levels for GM-CSF, IL-15, IL-10, IFNg, and CXCL9, from samples shown in (F). Summary data is presented as mean ± SEM. *p < 0.05, **p < 0.01 by ordinary one-way ANOVA with Dunnett’s multiple comparisons testing (calculated in GraphPad). Only showing significant results for comparisons between Tx and other conditions.

A close examination of all clusters revealed that 9 clusters described communication between myeloid cell types and 9 described communication between myeloid cells and T cells (Fig. S6C). All of these clusters were enriched for cell-cell communication axes that were supported by the literature (Table S1). Interestingly, despite previous work primarily implicating DCs in CD40ag-mediated anti-tumor immunity,^20,26,27,35,57^ only two of the 11 clusters enriched in the ICB lo + CD40ag-treated samples included DC-to-T cell pairs (Fig. S6C). Cluster 16 was primarily made up of cDC1-to-Treg pairs predicted to interact through the MHCII antigen presenting complex (Fig. S6D). Cluster 13 was primarily made up of mregDC-to-Treg pairs predicted to interact along pro-inflammatory axes, including *Il12b*–*Il12rb(1,2)* and *Il15*–*Il12rb*, as well as anti-inflammatory axes, including *Ccl22*–*Ccr4* (Fig. 5B). We further noted that cluster 15 comprised Treg-to-mregDC pairs predicted to signal through *Il10-Il10ra* and *Csf1-Csf1r* (Fig. 5B). We found a significant overlap in the mregDCs predicted to communicate with Tregs in clusters 13 and 15 (∼30%; Fig. 5C). We previously demonstrated that CD40ag-induced, DC-secreted IL-12 signaling can drive tumor regression in YR1.7 melanomas,^20^ but these results suggested that IL-12 could also be simultaneously amplifying immunosuppressive signals at this early time point. To directly test the role of IL-12 signaling in the antitumor response, we administered IL-12 blocking antibody (αIL-12) or isotype control (rat IgG2a) 6 days post-tumor inoculation in YR1.7-bearing mice. We then treated mice at day 7 with ICB lo + CD40ag and monitored their tumor volume over time. When regression was not achieved, ICB lo + CD40ag + αIL-12-treated tumors grew out faster than ICB lo + CD40ag + IgG2a-treated tumors, reaching endpoint (1 cm^3^ or ulceration) as early as 26 days post-tumor initiation (Fig. S7A). However, both treated cohorts, with and without IL-12 blockade, achieved the same overall survival of 60% (Fig. S7B). This result suggests that, while IL-12 contributes to the acute antitumor response we observe with ICB lo+CD40ag, there are additional therapy-induced mechanisms at play which contribute to sustaining tumor regression over time.

Our computational analyses also proposed ICB lo + CD40ag-specific axes of macrophage-T cell communication. Clusters 1 and 14 were comprised primarily of macrophage-to-CD8^+^ T cell and CD8^+^ T cell-to-macrophage pairs, respectively (Figs. 5B and S6C). Cell-cell pairs in cluster 1 were predicted to interact via *Cxcl9/10*–*Cxcr3* (Fig. 5B), axes that are required for CD8^+^ T cell infiltration into the TME and subsequent response to ICB,^51,58^ in addition to *Cxcl16*– *Cxcr6* and *Il18*–*Il18r1* (Fig. 5B), similar to CD40ag monotherapy. CXCL16-CXCR6 has been shown to aid in CD8^+^ T cell positioning in the TME by recruiting T cells to perivascular niches,^18^ but macrophages have not previously been shown to be the source of CXCL16. Pairs in cluster 14 were predicted to interact via IFNγ and GM-CSF (Fig. 5B), signals which can amplify pro-inflammatory myeloid activation.^59,60^ When we identified these cells in the gene expression space, we confirmed that many of the same T cells were both senders and receivers with macrophages (Fig. 5D), suggesting the possibility of a CD40ag-induced feedback loop.

Notably, CD40ag + ICB lo also decreased the frequency of a cluster of Treg-to-macrophage pairs enriched for IL-10 signaling (Cluster 9, Figs. 5B and S6C). When we mapped macrophages interacting with Tregs in the gene expression space (Cluster 9; Fig. 5D), we found that they were distinct from the macrophages interacting with CD4/CD8^+^ T cells (Clusters 1, 14; Fig. 5D). Experimental analysis of YR1.7 tumors from IL-10-GFP reporter mice treated with CD40ag + ICB lo revealed that the addition of CD40ag did not alter CD4^+^IL-10^+^ T density relative to ICB lo treatment alone (Fig. S5B). This was supported by Milo analysis on the scRNA-seq data, which did not identify a significant reduction in the frequency of Tregs following combinatorial therapy relative to ICB lo-treated tumors (Fig. S8). Instead, in the tissue samples, we observed a significant decrease in proximity (within 50μm and 100μm) between macrophages and immunosuppressive T cells (CD4^+^IL-10^+^) in the TME with combinatorial treatment relative to ICB lo and control (Fig. S5C-D). This result suggested that including CD40ag in the treatment regimen might instead rewire this axis by altering macrophage phenotype and spatial orientation to break immunosuppressive signaling.

Our analysis implicated both IL-12 signaling via DCs and IL-18 signaling via macrophages in the acute response to ICB lo + CD40ag (Fig. 5B-D). We therefore tested how blocking IL-12 and IL-18 would affect systemic response to treatment by analyzing acute changes in cytokine/chemokine (C/C) networks induced with therapy. To do this, we collected serum 24 hours post-therapy from ICB lo + CD40ag-treated mice, which had been treated with αIL-12 and/or IL18BP or its respective control, and conducted systemic cytokine profiling (Fig. 5E). Hierarchical clustering identified three major modules of C/C activation that were differentially affected by the blockade of IL-12 and IL-18 (Fig. 5F). Module 1 included treatment-induced C/Cs whose secretion was decreased by blocking IL-12, IL-18, or both (Fig. 5F; ‘IL-12- and IL-18-dependent’, in blue). C/Cs in Module 1 included GM-CSF, IL-15, IL-10, IL-12p40, and M-CSF (Figs. 5G and S9). This is consistent with our hypothesis that ICB lo + CD40ag activates complex combinatorial networks that are not driven solely by IL-12. However, blocking IL-12 did more strongly modulate some signals, including IFNγ and TNF (Figs. 5F-G and S9; ‘IL-12-dependent’, in yellow). Finally, secretion of C/Cs in Module 3 (Fig. 5F; ‘independent’, in turquoise), which included CXCL9 (Fig. 5G), were not affected by blocking IL-12 and IL-18 separately or in combination. As we predicted ICB lo + CD40ag to uniquely upregulate macrophage-T cell crosstalk via CXCL9 compared to control and ICB alone (Fig. 5B, Cluster 1), this result suggests that CXCL9 could constitute an additional axis of treatment-induced cell-cell communication which is independent of IL-12/IL-18 and contributes to emergent behaviors in the TME. Overall, these results suggest that macrophages and DCs, via IL-18 and IL-12, respectively, play distinct antitumor roles in the complex signaling networks activated by treatment with ICB lo + CD40ag.

### Spatial transcriptomics analysis validates therapy-induced changes in intratumoral macrophage-T cell subset interaction predicted by single-cell-resolved interactomes

Our scRNA-seq analyses show that resolving cell-cell communication at single-cell resolution can identify macrophages with distinct signaling partners that correspond to distinct transcriptomic states (Fig. 5C-D). This suggests that our method might reveal features of local neighborhood interactions despite the fact that scRNA-seq data is not spatially resolved. To determine if the macrophages that we predict to participate in different cell-cell communication clusters corresponded to their spatial adjacency with other cell types, we collected untreated, ICB lo-, and ICB lo+CD40ag-treated tumors 24 hours post-treatment for spatial transcriptomics analysis via deterministic barcoding in tissue for spatial omics sequencing (DBiT-seq).^61^ We identified gene signatures for macrophages, mregDCs, Tregs, and CD8^+^ T cells from our scRNA-seq data which we used to identify the presence of these cells in the DBiT-seq spots (Figs. 6A and S10A).

**Figure 6.**
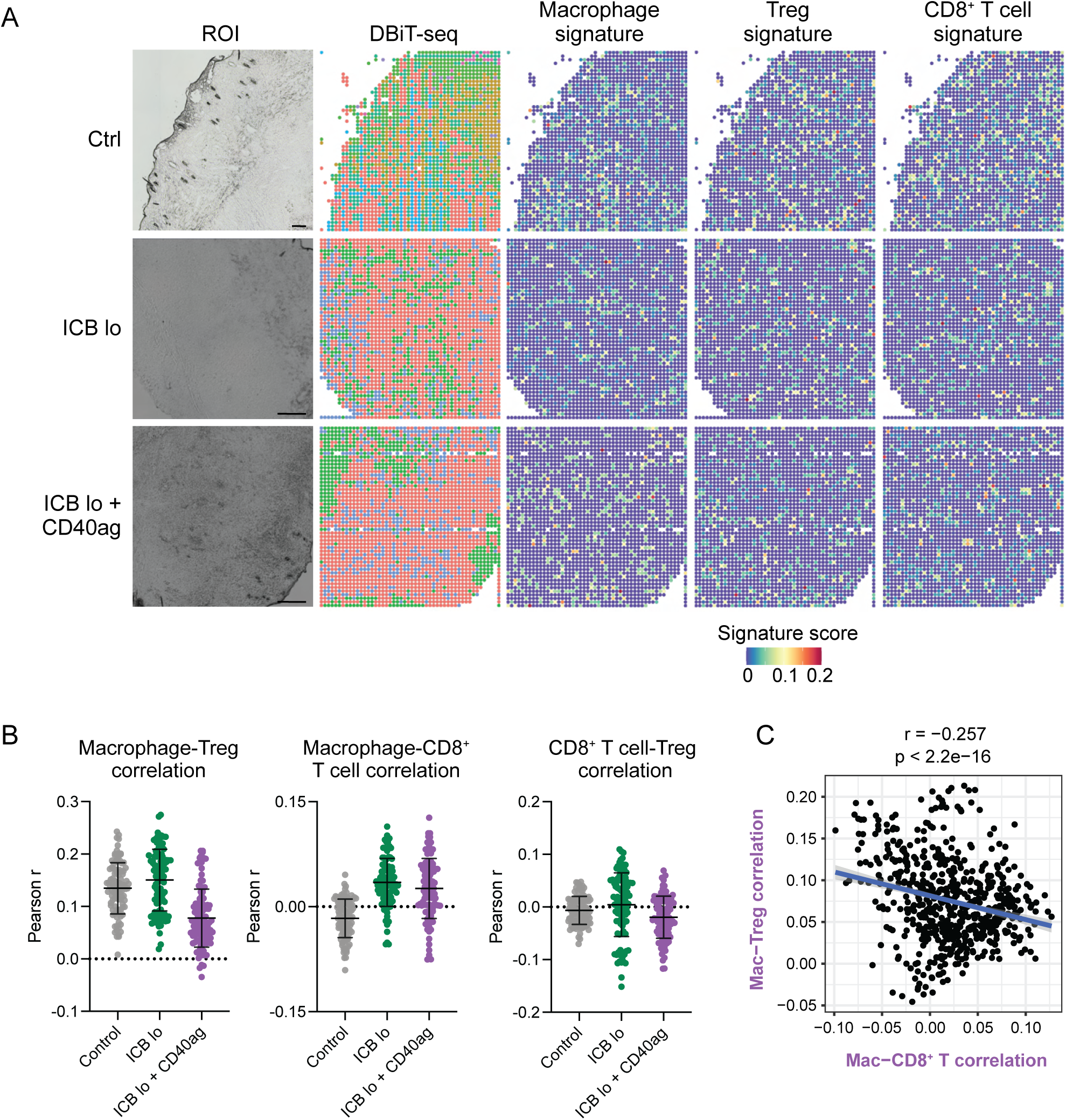
ICB and CD40ag acutely and differentially alter immune spatial orientation and inflammatory signaling. (A) DBiT-seq spatial transcriptomics analysis of YR1.7 tumors harvested from mice 24 hours post-treatment where applicable. Left: Region of interest (ROI). Middle: DBiT-seq spots. Right: macrophage, Treg, and CD8^+^ T cell signature scores for each pixel in the spatial transcriptomics spot map demonstrating the presence of each cell type in the TME. Scale bars are 200 μm. (B) Scatter plots demonstrating per-pixel Pearson correlation coefficients between (left) macrophage and Treg signature scores, (middle) macrophage and CD8^+^ T cell signature scores, and (right) CD8 ^+^ T cell and Treg signature scores for 100 20×20 grids randomly sampled from each DBiT-seq sample (control – grey, ICB lo – green, ICB lo + CD40ag – purple). Summary data is presented as mean ± SD. (C) Scatter plot comparing correlation between macrophage and CD8^+^ T cell signature scores (x-axis) to correlation between macrophage and Treg signature scores (y-axis) for corresponding 1000 20×20 grids randomly sampled from the ICB lo + CD40ag-treated DBiT-seq sample.

Our previous analyses suggested that combining CD40ag with ICB lo rewired macrophages away from immunosuppressive interactions with Tregs and toward productive interactions with CD8^+^ T cells. To provide evidence to support this hypothesis, we analyzed how different treatments changed the spatial adjacency of macrophages and T cell subsets. We found that, compared to control, treatment with ICB lo alone and ICB lo + CD40ag increased macrophage-CD8^+^ T cell correlation within 24 hours of administration (Fig. 6B, center). In contrast, compared to control, only the combination treatment decreased macrophage-Treg correlation, while ICB lo yielded a slight increase (Fig. 6B, left). In comparison, while mregDC-Treg correlation followed a similar trend as macrophage-Treg correlation across conditions, we could not assert an increase in mregDC-CD8^+^ T cell correlation with treatment within 24 hours of administration (Fig. S10B). Importantly, directly comparing macrophage-CD8^+^ T cell correlation to macrophage-Treg correlation within the ICB lo + CD40ag-treated sample revealed an inverse relationship (Fig. 6C), suggesting that distinct macrophages engage with different T cell subsets in the TME. Altogether, our spatial analyses support our hypothesis that combinatorial treatment with CD40ag and ICB decreases signaling between macrophages and Tregs while simultaneously increasing signaling between macrophages and CD8^+^ T cells to initiate antitumor immunity-promoting myeloid-T cell crosstalk that ultimately supports tumor regression.

### Cell-cell interactions predicted to be activated by CD40ag+ICB and maintained over time suggest the presence of immune hubs

Finally, we wanted to understand how antitumor immunity-promoting myeloid-T cell crosstalk persisted over time. To do this, we collected scRNA-seq data from tumors treated with ICB lo + CD40ag at 3 days post-treatment (10 days post-tumor initiation) and compared it to scRNA-seq data from tumors collected 24 hours post-therapy. We generated NICHES networks for treated samples across post-therapy timepoints and found that nearly two-thirds (239/363) of Milo-assigned neighborhoods were differentially abundant w.r.t. time post-treatment (Fig. 7A-B), indicating many changes in the landscape of predicted cell-cell communication over time.

**Figure 7.**
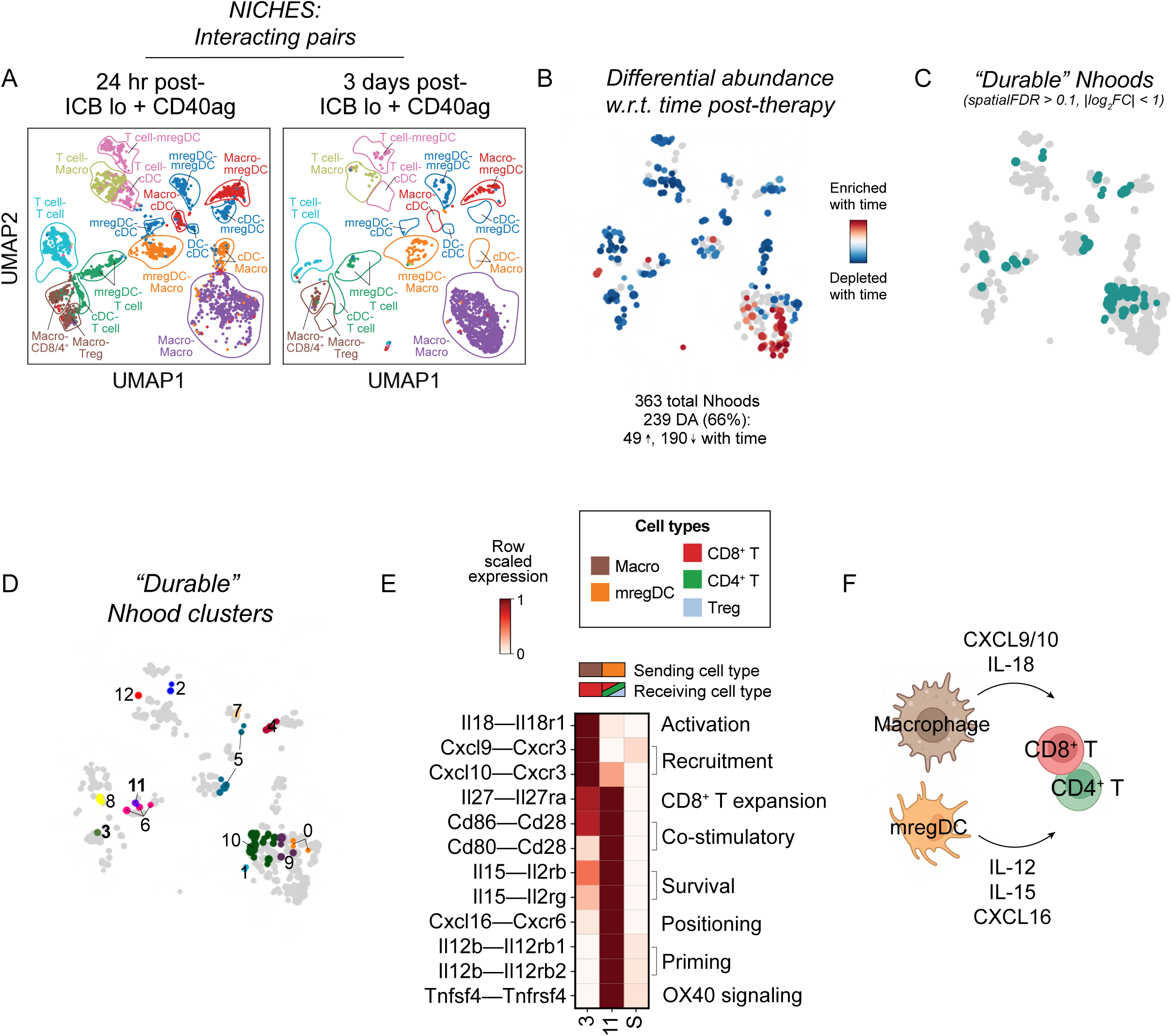
Characterization of ‘durable’ therapy-induced interactions across time. (A) UMAP embedding of NICHES interactomes for ICB lo+CD40ag-treated samples collected at 24- and 72-hours post-treatment separated by timepoint and colored by broad cell type pair. (B) UMAP embedding of 363 Milo-assigned ‘neighborhoods’ (nhoods) of cell-cell pairs colored by differential abundance result w.r.t. time post-treatment. Grey nhoods were not differentially abundant (n=124), while blue (n=190) and red (n=49) nhoods decreased and increased in frequency, respectively, with time post-treatment. (C-D) UMAP embedding as in (B) colored by ‘durable’ nhoods (spatialFDR > 0.1, abs(log_2_FC) < 1), which were grouped into 13 clusters for downstream analysis. (E) Heatmap depicting per-row scaled interaction scores for selected ligand-receptor axes (y-axis) across ‘durable’ clusters 3 and 11 and compared to all differentially abundant neighborhoods (i.e., S) (x-axis). The key above the heatmap distinguishes which cell type sender-receiver pairs are involved in each highlighted cluster. (F) Schematic illustrating myeloid-T cell communication predicted to be durably upregulated with ICB lo + CD40ag.

To focus on combinatorial treatment-induced cell-cell interactions that are durable across time, we identified non-differentially abundant neighborhoods (Fig. 7C; spatial FDR > 0.1, |log_2_FC| < 1) and grouped them into 13 clusters for downstream analyses (i.e., ‘durable’ clusters, Fig. 7D). Clusters 3, composed of macrophage-to-CD8^+^ T cell interactions, was enriched for communication along *Il18*–*Il18r1* and *Cxcl9/10*–*Cxcr3* (Fig. 7E), the same T cell-recruiting and activating axes observed previously (Fig. 5B, Cluster 1). In addition, we identified Cluster 11, composed of mregDC-to-mixed T cell pairs, that was enriched for communication via IL-12 and IL-15 (Fig. 7E), similar to the mregDC-to-Treg cluster that emerged 24 hours post combination therapy (Cluster 13, Fig. 5B), although the composition of receiving T cell subsets was slightly altered. Importantly, in addition to IL-12 and IL-15 signaling, this cluster also included *Tnfsf4* (OX40L)–*Tnfrsf4* (OX40), *Cd80/86*–*Cd28*, and *Cxcl16*–*Cxcr6* interactions (Fig. 7E), mimicking the signature of mregDCs (i.e., cDC3s) involved in CD8^+^ T cell positioning in the TME at perivascular niches.^18^ Collectively, these results suggest that ICB and CD40ag combine to facilitate durable myeloid-T cell crosstalk via CD40ag-stimulated axes of cell-cell communication (Fig. 7F), including axes known to organize perivascular immunity hubs,^18^ ultimately driving antitumor immunity.

## DISCUSSION

Cancer immunotherapies that stimulate innate versus adaptive immune compartments are increasingly being combined in situations of ICB resistance because communication between these compartments is critical for restoring antitumor immunity. Here, we developed and validated a method to analyze scRNA-seq data to explore how innate-adaptive immune cell communication changes when myeloid-targeted CD40 agonist is combined with checkpoint inhibitors against PD-1 and CTLA-4. We found that CD40ag and ICB rapidly altered intratumoral immune composition and produced distinct effects on different myeloid cell subsets. These treatments ultimately rewired innate-adaptive immune crosstalk in the tumor that enabled tumor regression.

The key to discovering the underlying cell-cell signaling axes that mediated the CD40ag + ICB-driven antitumor response was the use of NICHES to construct scRNA-seq-derived interactomes at single-cell resolution.^10^ In particular, we demonstrated that macrophage gene expression becomes increasingly skewed after treatment with CD40ag, and so averaging across this population, as most cell-cell inference methods require,^4^ would have negated the highly granular profiling afforded by scRNA-seq. Moreover, sub-clustering these cells might have inappropriately enforced discretization of a rapidly evolving population of cells in flux between a variety of cell states. By preserving heterogeneity when inferring myeloid-T cell communication from scRNA-seq, we uncovered new axes of communication induced by CD40ag that were observed in multiple preclinical models (Fig. S3).

Further, using Milo for differential abundance testing allowed us to efficiently compare interactomes and generate hypotheses about treatment regimen-specific cell-cell communication axes. Unlike similar differential abundance testing algorithms,^62,63^ Milo’s statistical framework maintains greater flexibility, allowing for the accounting of replicates and for modeling more complex experimental designs,^41^ such as continuous covariates like treatment dose. The use of Milo is limited when comparing across datasets without replicates, as we demonstrated with GSE224400,^25^ and as is often the case due to the high cost per scRNA-seq sample. However, recent advances in sample multiplexing and sequencing instrumentation will continue to lower this barrier and support the use of Milo to efficiently prune NICHES-inferred cell-cell communication networks with greater statistical power.

By efficiently handling myeloid cell heterogeneity in scRNA-seq data, our study provided new insights into how CD40ag rewires cell-cell communication in the tumor. CD40ag monotherapy has been met with mixed success in the clinic.^57^ In line with this, we observed that although CD40ag monotherapy slows tumor growth in YR1.7-bearing mice, this trend is not sustained over time (Fig. 1B-D). Our analysis of CD40ag-induced cell-cell communication suggested some potential mechanisms of resistance to monotherapy across tumor models, namely myeloid-T cell crosstalk via IL-10, PD-(L)1, and CCL22 (Figs. 3D-E and S3E-F). While mregDCs are known to be CD40ag-inducible^20,27^ and are typically thought to support antitumor immunity,^18,42^ recent work has implicated a suppressive Treg-mregDC-lymphatic niche in tumor stroma, formed via CCL22-CCR4-driven chemotaxis, in impeding antitumor adaptive immune activation.^45^ We hypothesize that, with CD40ag monotherapy, the balance between antitumor and immunosuppressive myeloid-driven T cell activation might flip over time, with inflammatory signaling driving tumor regression in the short-term, while also amplifying communication that restrains the antitumor adaptive immune response in the long-term. Further study is required to understand the cell-cell communication which underlies this non-monotonic trend in tumor growth that we observe over time following treatment with CD40ag monotherapy.

Importantly, changes in cell-cell communication induced by CD40ag are sustained in the presence of ICB. While CD40ag induced mregDC-Treg communication in both contexts (Fig. 5B, compare to Fig. 3D-E), ICB lo + CD40ag treatment additionally induced a macrophage–CD8^+^ T cell signaling loop via IL-18, CXCL9/10, and CXCL16 (from macrophages to T cells), and IFNγ and GM-CSF (from T cells back to macrophages). We hypothesize that this is driven by a macrophage subset that emerges exclusively with combinatorial therapy. Macrophage-derived CXCL9 and CXCL10 have previously been described to be required for response to ICB.^51^ Interestingly, although CXCL16–CXCR6 signaling has been shown to support antitumor immunity by promoting the survival and expansion of CD8^+^ T cells in the TME,^18^ DCs and not macrophages are typically thought to mediate this interaction.^64^ However, a study in sepsis linked CXCL16-expressing macrophages to the accumulation of antitumoral T cells in tissues, including a subset of CD8^+^ T cells,^65^ similar to our results. Interestingly, by 72-hours post-therapy, we found that *Cxcl16*–*Cxcr6* signaling was enriched in a mregDC-to-mixed T cell cluster (Fig. 7E), suggested that the two myeloid cell types might cooperate to prime and effect antitumor immune responses. We also identified an ICB lo-specific Treg-to-macrophage signaling cluster enriched for IL-10 that was reduced by both increasing ICB dose (Fig. 4D-F, ‘Cluster 8’) and by adding CD40ag (Fig. 5A-B, ‘Cluster 9’). Interestingly, increasing ICB dose depleted Tregs, which we confirmed with immunofluorescence (Figs. 4H-J and S5B). In contrast, we found that adding CD40ag did not acutely deplete IL-10^+^CD4^+^ cell density in the TME relative to ICB lo treatment; rather, we observed a decrease in the fraction of macrophages within a ‘communication radius’ of IL-10^+^CD4^+^ cells with ICB lo + CD40ag treatment that we did not observe with increased ICB dose. Thus, while our computational analyses demonstrated that both increasing ICB dose and adding CD40ag decreased Treg immunosuppressive signaling via IL-10, we uncovered different mechanisms for achieving this outcome. While increasing ICB dose breaks immunosuppressive IL-10–IL10R Treg–TAM signaling via Treg depletion, CD40ag appears to change TAM state, restricting their communication and/or adjacency in the tumor.

To provide experimental support for our hypotheses, we spatially profiled tumor samples across treatments with DBiT-seq.^61^ We found that our predicted changes in macrophage–T cell communication were supported by expected changes in adjacency in the tumor (Fig. 6B-C). However, the resolution of the DBiT-seq data was not sufficient to confirm specific ligand-receptor pairing. It is also important to note that, despite DCs being previously implicated to drive CD8^+^ T cell activation in response to CD40ag,^20,27,35^ we did not observe an increase in mregDC-CD8^+^ T cell adjacency relative to control 24 hours post-combinatorial therapy. Interestingly, treatment with ICB lo + CD40ag also yielded the greatest variability in correlation between macrophage and CD8^+^ T cell signatures among subsampled partitions of the tumor slice (Fig. 6B, center). Although this could be connected to many biological and technical factors, one hypothesis is that, in different sections of the same tumor, treatment with ICB lo and CD40ag yields regions of immunosuppression and tumor regression. This phenomenon has been previously observed in patients with cutaneous melanoma.^66^ Further work is required to more completely understand the spatial heterogeneity in myeloid-T cell signaling that is captured in preclinical models of melanoma.

Finally, many of our analyses revealed more differentially abundant clusters than those that were highlighted (e.g., Figs. 5A, S6, and Table S1). Of the 11 ICB lo + CD40ag-specific clusters we identified, 7 involved DC subsets, 2 of which specifically predicted mregDC-to-mregDC communication. Given that we know DCs comprise <5% of total intratumoral immune cells in our tumor model (Fig. S1C), it is unlikely that they are encountering each other in the tumor, although autocrine signaling is possible. While enriching for DCs prior to sequencing allowed us to understand how different DC subsets responded to immunotherapy, this may have also artificially inflated the likelihood of DC-involved cell-cell communication in our analyses.

Moreover, we narrowed our focus to crosstalk between cell types directly downstream of treatment with ICB and CD40ag to only include T cell subsets, macrophages and DCs. Indeed, neutrophils^25^ and B cells^67,68^ have both been shown to be CD40ag-responsive in other contexts, although previous work by our group has suggested that CD40ag does not require neutrophils to be efficacious in models of melanoma.^21^ Further, B cell content in the TME has been shown to not be associated with response to anti-PD-1 or overall survival, both in patients with melanoma and in YR1.7-bearing mice,^69^ nor is B cell infiltration into the TME largely altered by treatment with CD40ag (Krykbaeva et al.^20^ and Fig. S1C). Future work is necessary to definitively elucidate whether these cell types participate in tumor regression-promoting interactions in response to combinatorial therapy with ICB and CD40ag.

## RESOURCE AVAILABILITY

### Lead contact

Further information and requests for resources and reagents should be directed to and will be fulfilled by the lead contact, Kathryn Miller-Jensen.

### Materials availability

This study did not generate new unique reagents.

### Data and code availability

- The scRNA-seq data generated in this study are publicly available in Gene Expression Omnibus (GEO) at GSEXXXXXX (to be provided upon publication). The scRNA-seq data describing 9-day CD40ag-treated MC38 tumors (Fig. S3) are publicly available in GEO at GSE224400.
- The code used to generate the figures presented in this study is available in the following public GitHub repository: [https://github.com/khbridges/Bridgesetal-CCC].
- All other data supporting the findings of this study are available from the corresponding author upon reasonable request.

## ACKNOWLEDGMENTS

This work was supported by the NIH/NCI U01CA238728 award (to K. Miller-Jensen and M. Bosenberg). The authors thank Goran Micevic, Andrew Daniels, Meaghan McGeary, Ronan Talty, and the Yale Center for Genome Analysis (YCGA) for assistance with scRNA-seq data collection. The authors further thank all members of the Miller-Jensen and Kaech labs for insightful discussions and advice.

## AUTHOR CONTRIBUTIONS

Conceptualization, K.B., G.A.P., W.D., M.W.B., R.F., S.M.K., and K.M-J.; Software, K.B., A.B., and J.P.B.; Formal analysis, K.B., A.B., and J.P.B.; Investigation, G.A.P, W.D., A.B., J.P.B., Z.X., V.M., V.T., M.L., and K.P.; Resources, M.W.B., R.F., S.M.K., and K.M-J. Data Curation, K.B. and A.B.; Writing – Original Draft, K.B. and K.M-J.; Writing – Review & Editing, K.B., G.A.P., and K.M-J.; Visualization, K.B. and K.M-J.; Supervision, H.K., M.W.B., R.F., S.M.K., and K.M-J. Funding Acquisition, M.W.B. and K.M-J.

## STAR METHODS

### KEY RESOURCES TABLE

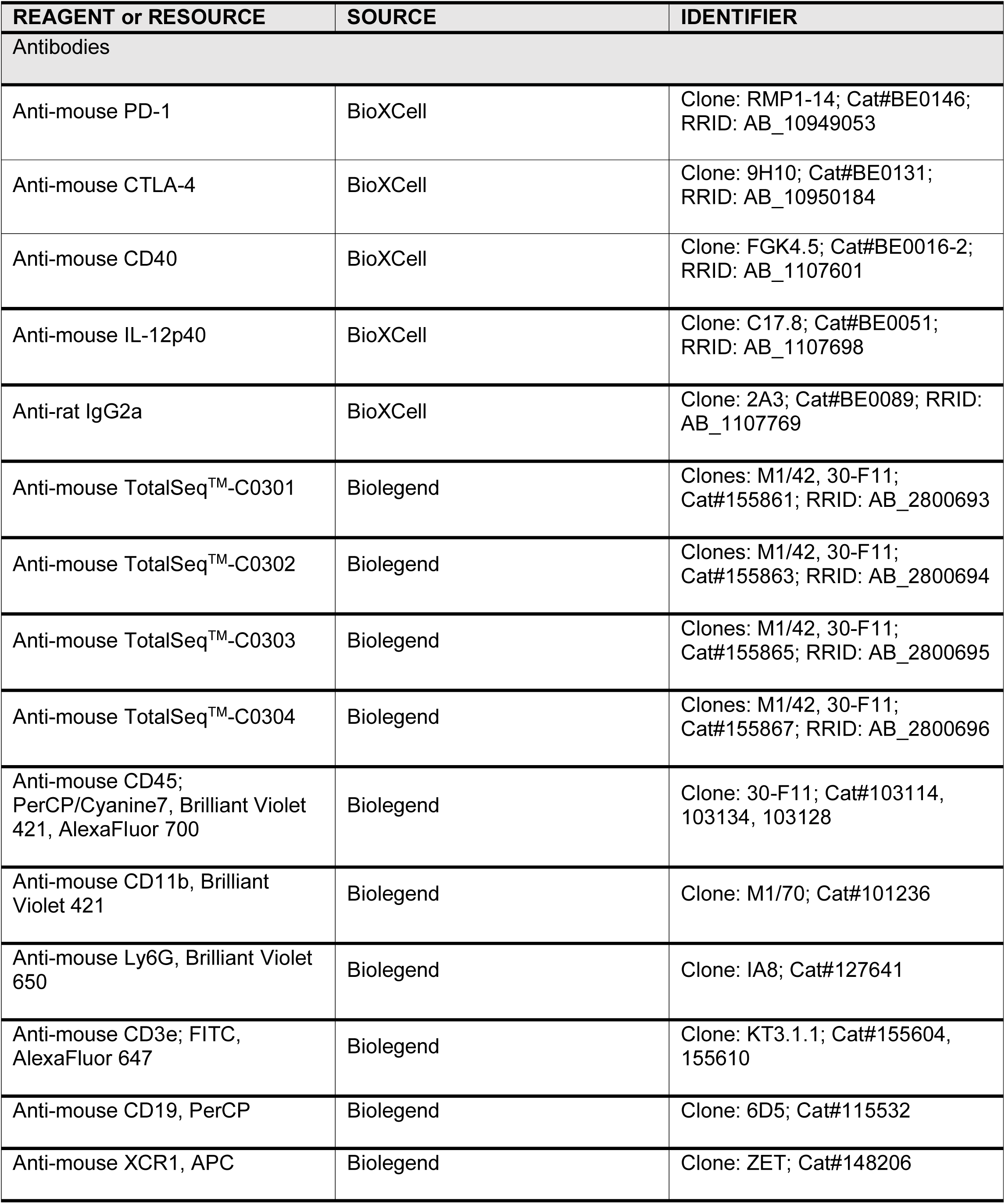
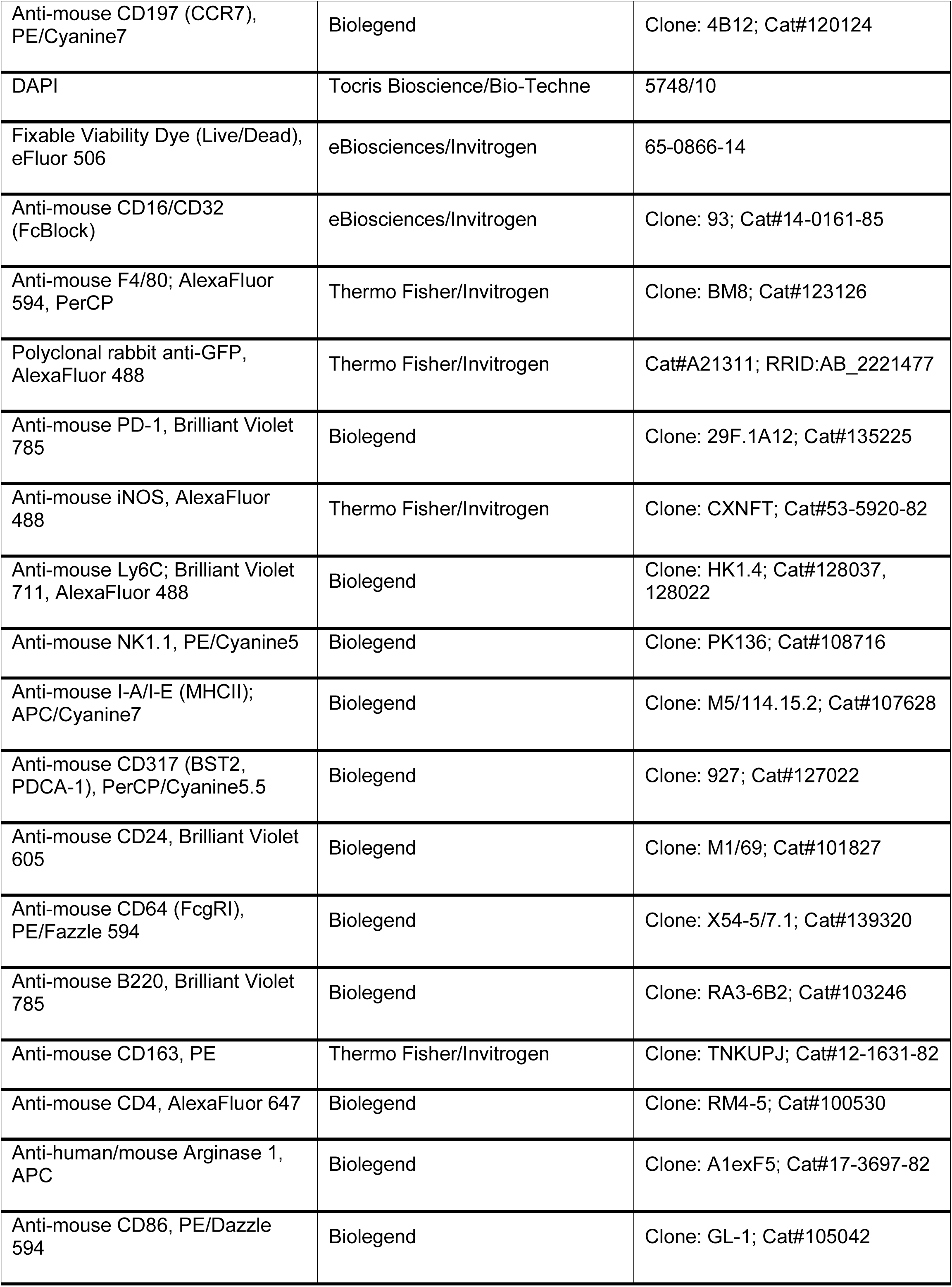
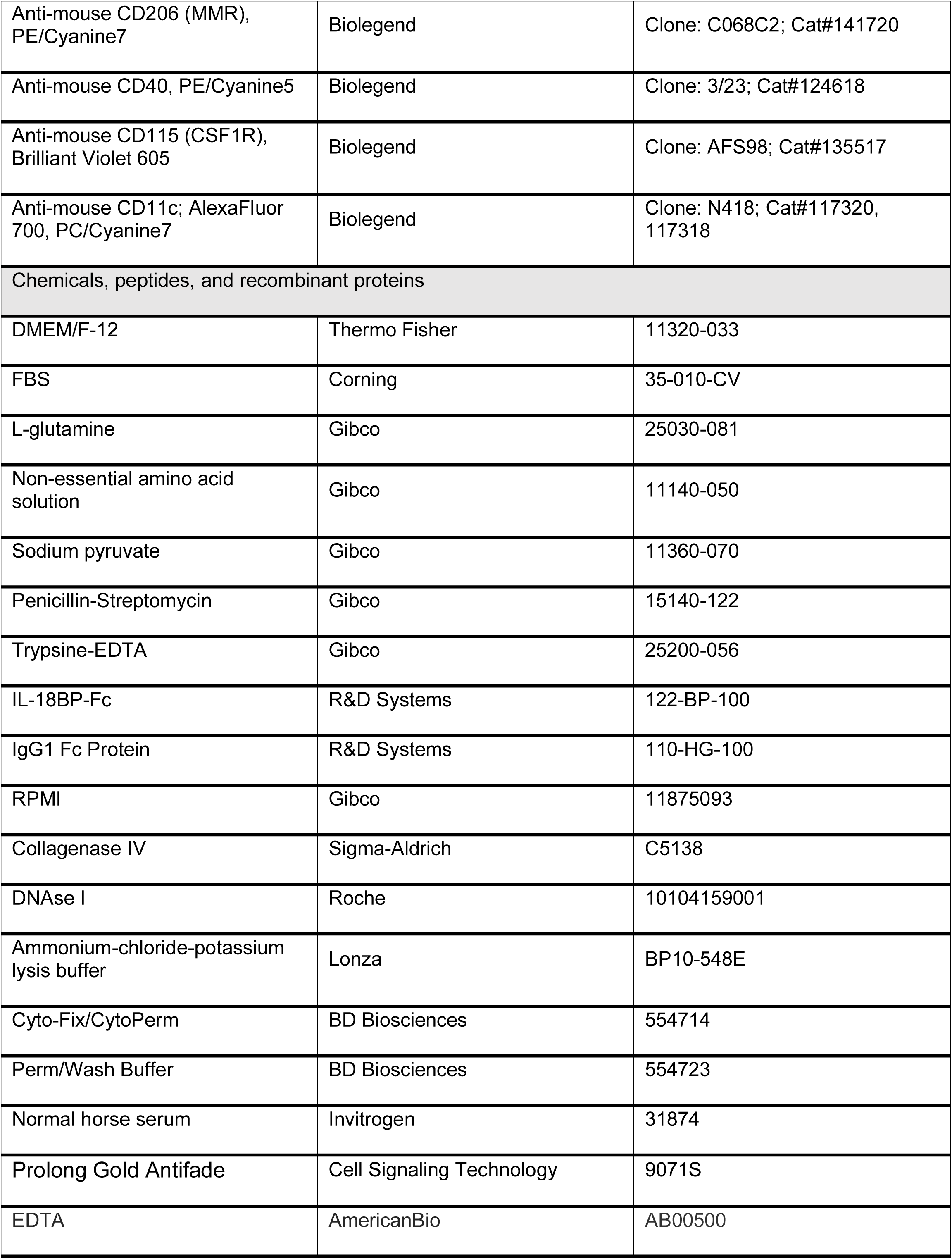
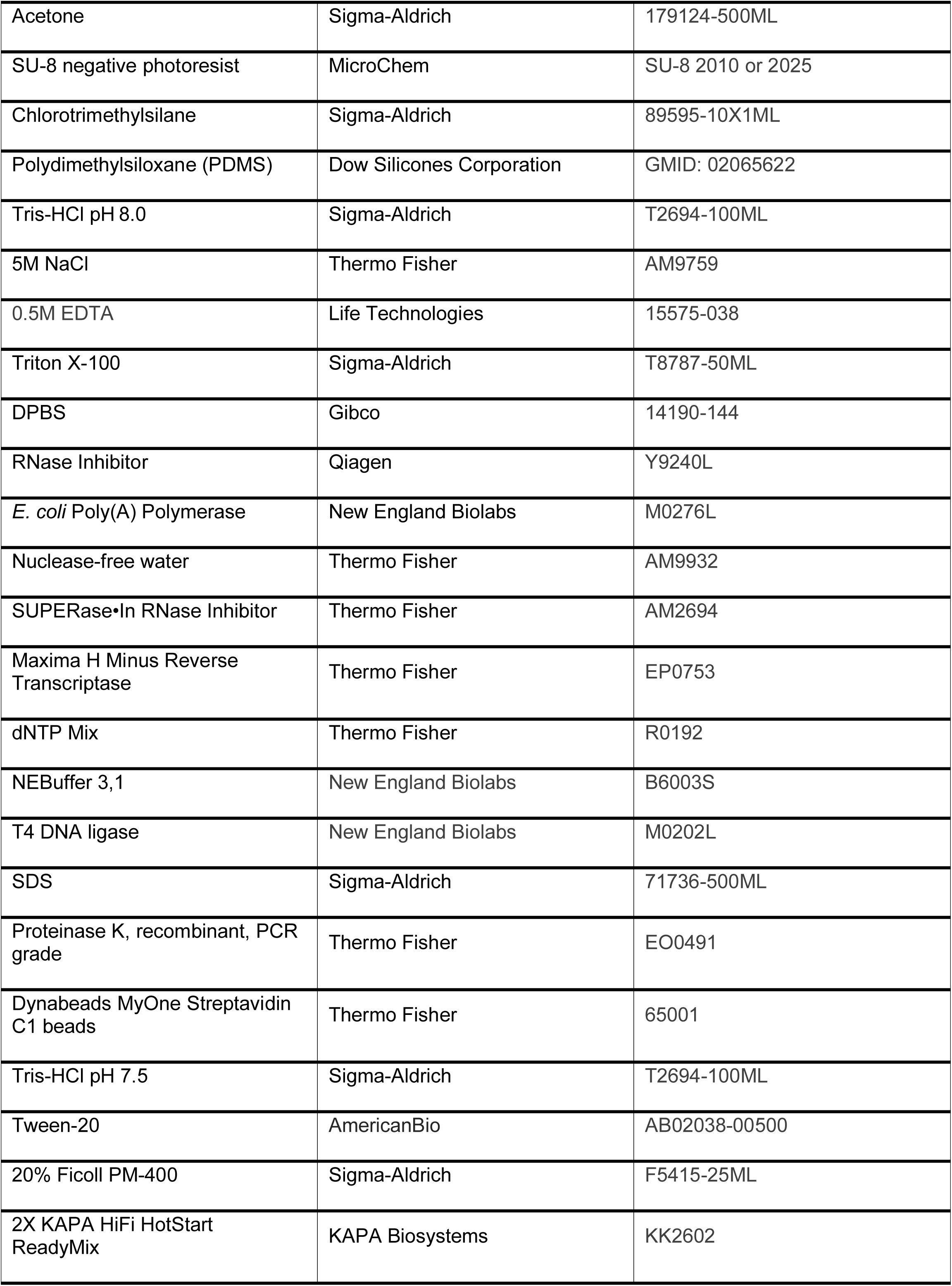
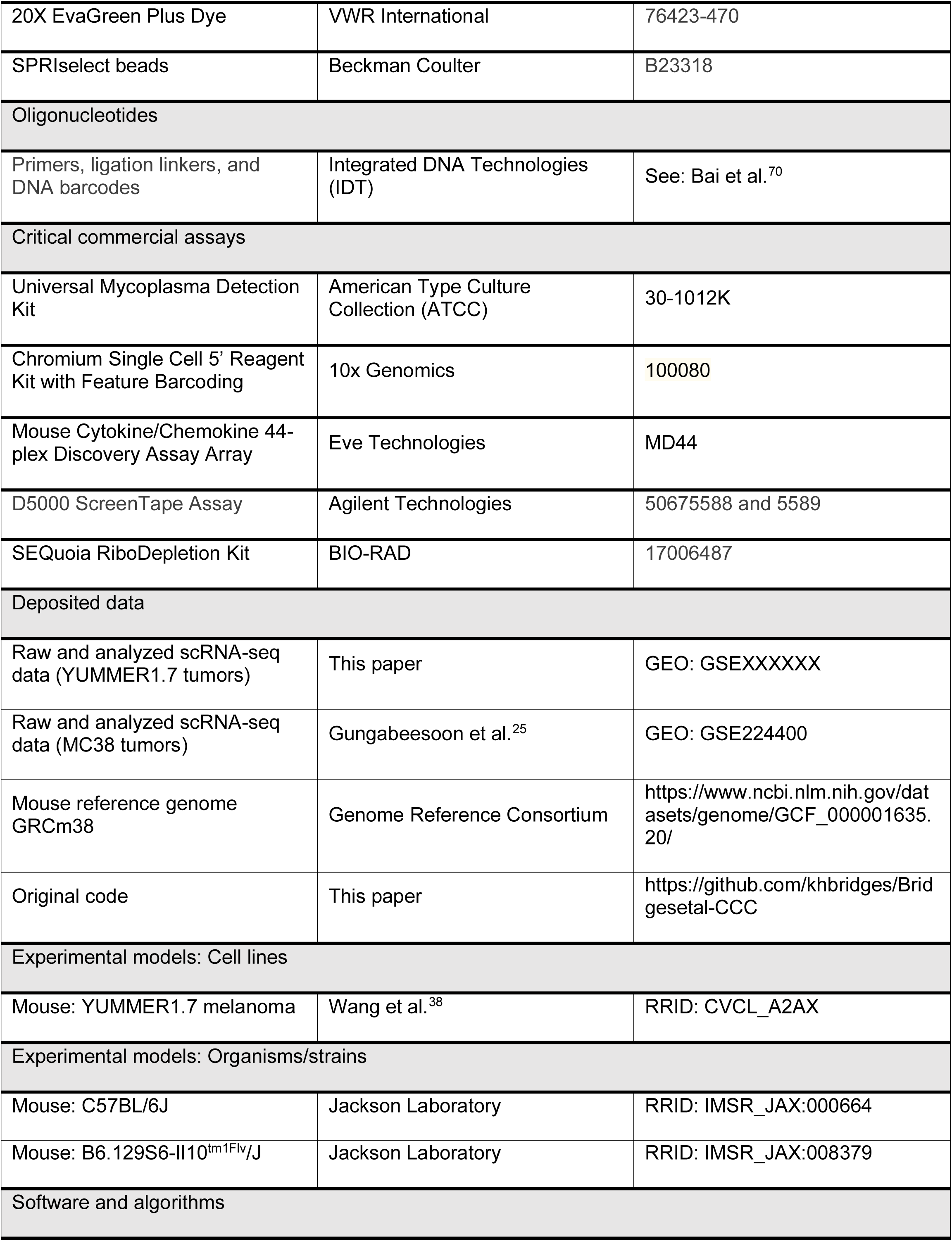
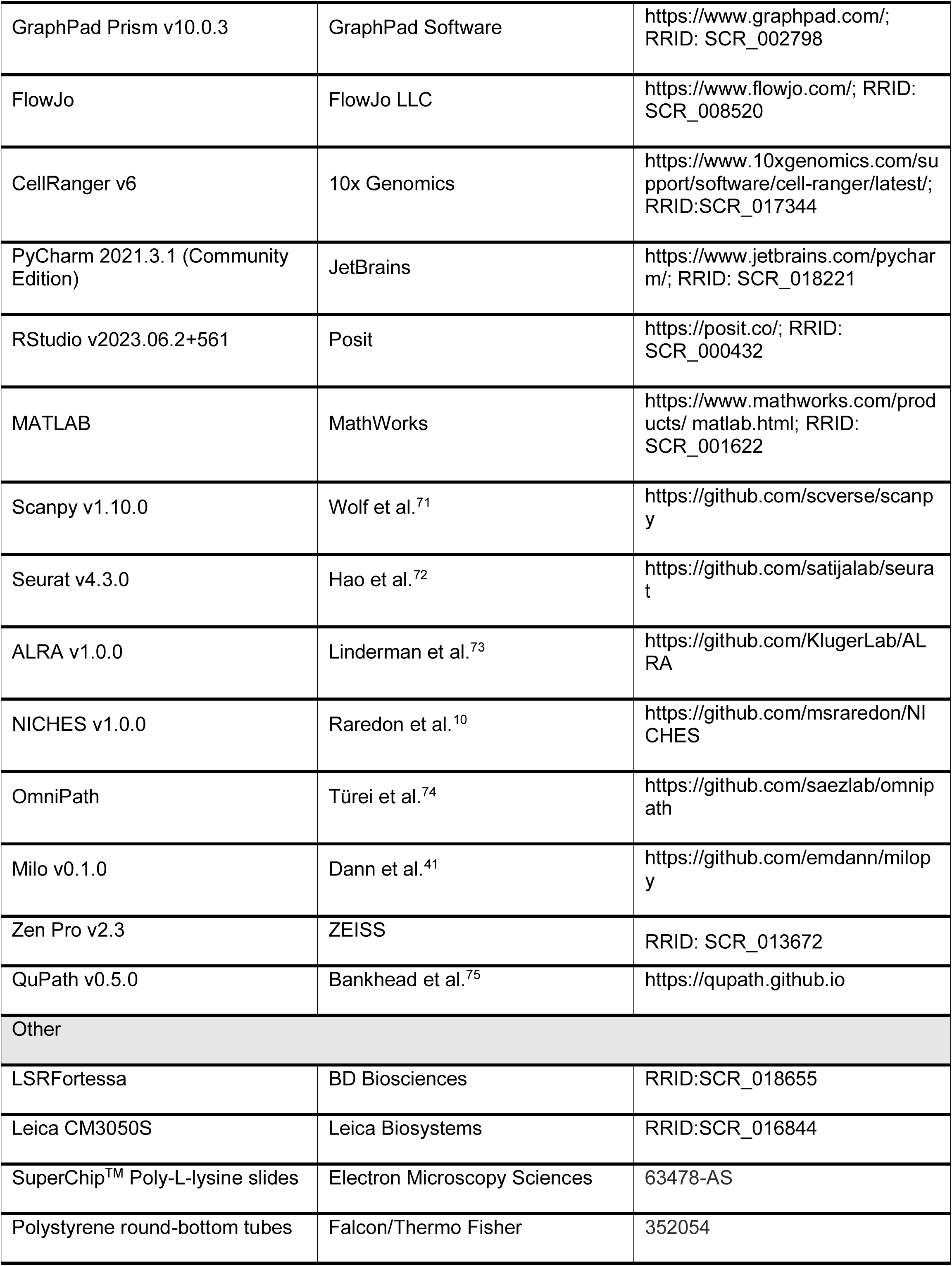
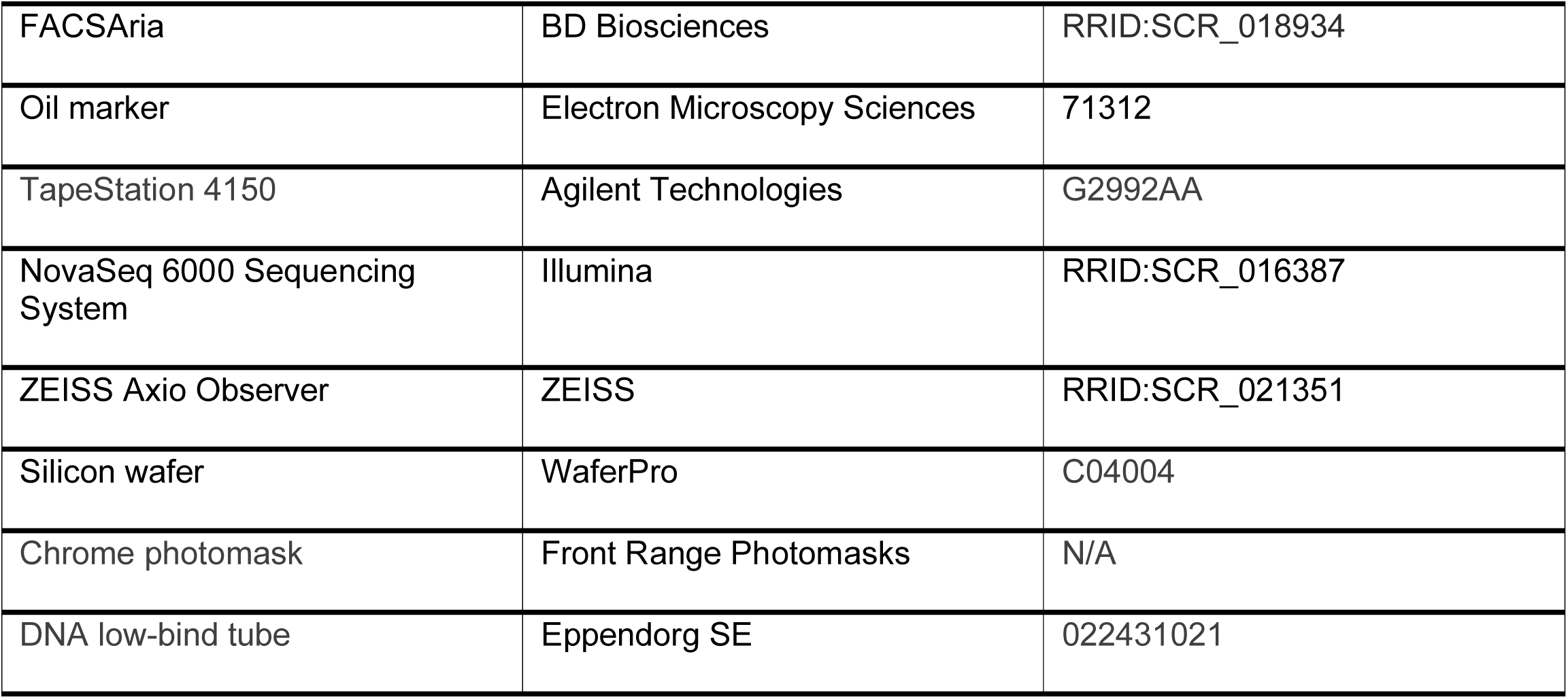

## EXPERIMENTAL MODEL AND SUBJECT DETAILS

### Mice

Wild type (WT) C57BL/6J mice and B6.129S6-Il10^tm1Flv^/J (IL-10-GFP reporter) mice were purchased from Jackson Laboratories. Mice were housed and maintained according to the standard pathogen-free conditions of the Yale Animal Resources Center facilities. All animal studies were conducted in accordance with protocols approved by the Yale University Institutional Animal Care and Use Committee (IACUC).

### Cell lines

YUMMER1.7 (YR1.7) cells^38^ were cultured in DMEM/F-12 and supplemented with 10% FBS, 2mM L-glutamine, 1% non-essential amino acids, 1% sodium pyruvate, and 1% penicillin and streptomycin. All cells were incubated at 37°C and 5% CO_2_. Cells were cultured in tissue culture-treated plastic flasks, detached using 0.25% Trypsine-EDTA, and kept at low passage (less than 10), showing negative test results for mycoplasma contamination within the last year.

## METHOD DETAILS

### Tumor induction and *in vivo* treatments

Mice were anesthetized with isoflurane and injected subcutaneously (s.c.) with 0.5×10^6^ YR1.7 cells on each flank. Tumors which displayed expected growth (i.e., 100-200 mm^3^ at 7 days post-tumor inoculation) were included in the studies. Where applicable, mice were treated with blocking antibodies against PD-1 (αPD-1) at 50μg/mouse (low dose) or 200μg/mouse (high dose) and against CTLA-4 (αCTLA-4) at 50μg/mouse (low dose) or 200μg/mouse (high dose), and/or with agonistic antibody against CD40 (CD40ag) at 50μg/mouse. For survival experiments, treatments were administered by intraperitoneal (i.p.) injection beginning at 7-days post-tumor inoculation. Treatments were repeated every 3 days for a total of five treatments. For cytokine depletion experiments, mice were additionally treated by i.p. injection with 500μg αIL-12 and/or 10μg IL-18BP-Fc, or with their respective isotype controls (500μg rat IgG2a and/or 10μg IgG1 Fc Protein) at 6-days post-tumor inoculation (24 hours prior to the initiation of treatment). When tumor growth and survival was assessed, a second depletion was administered at 13-days post-inoculation. Tumors were measured by calipers every 3-4 days. Tumor volume was estimated using the formula 0.5 x [(width)^2^ x length]. Humane endpoint was determined when tumors reached >1cm^3^, or when mice displayed deteriorating health, as per Yale University IACUC approved protocols.

### Tumor processing

At the designated timepoints, mice were euthanized in a CO_2_ chamber according to approved protocols, and tumors were resected and weighed for processing, as previously described.^20,76^ For single-cell suspensions, tumor tissues were minced and then incubated in digestion buffer (RPMI containing 2% FBS, 1 mg/mL collagenase IV and 0.1 mg/mL DNAse I) with agitation at 37°C for 30 minutes. Samples were kept on ice and filtered through a 70μm strainer. Red blood cells were lysed with ammonium-chloride-potassium (ACK) lysis buffer at room temperature (RT) for 5 min, and single-cell suspensions were washed and resuspended for counting and subsequent analysis. For tissue cryosectioning, tumors were cut in half and fixed in 2% paraformaldehyde (PFA) in PBS overnight at 4°C. Samples were then washed twice in PBS and transferred into a 30% sucrose in PBS solution for an additional overnight incubation at 4°C. Tumors were finally mounted in optimal cutting temperature (OCT) compound and preserved at - 80°C. OCT blocks were sectioned using a Leica CM3050S cryostat in 10μm tissue sections, placed onto positively charged slides, and further preserved at −80°C. For DBiT-seq, minor modifications were introduced to the tissue cryosectioning protocol: tumors were fixed in 0.1% PFA, and cryosections were mounted in poly-L-lysine-coated slides as previously described.^77^

### Immunostainings, flow cytometry, and fluorescence-activated cell sorting (FACS)

For single-cell suspensions prepared for flow cytometry assays, 1.0-5.0 x 10^6^ cells were transferred to polystyrene round-bottom tubes. Antibodies were pre-mixed according to each specific panel; extracellular mixes were prepared in FACS Buffer and intracellular mixes were prepared in 1X Perm/Wash Buffer. Cells were washed in FACS Buffer (2% FBS in PBS) by spinning at 400*g* for 5 min, resuspended, and then incubated with FcBlock 1/200 for 20 min at 4°C. Cells were then washed and incubated with the extracellular antibody mix for 30 min at 4°C. After washing again in FACS buffer, cells were fixed and permeabilized with Cyto-Fix/CytoPerm according to manufacturer instructions. Cells were then washed in Perm/Washer Buffer, the intracellular antibody mix was added, and incubated again for 30 min at 4°C. Cells were washed in FACS Buffer a final time, resuspended in PBS, and filtered through a 40μm strainer for analysis in an LSRFortessa. Flow cytometry data were processed in FlowJo. For single-cell suspensions prepared for CITE-seq (see below), cells from individual tumors were labeled with one of four TotalSeq^TM^-C anti-mouse hashtag antibodies (1μg per sample; see Key resources table) for 30 min at 4°C. Samples were then washed, and replicates for each experimental condition were combined. Cells were then stained for FACS sorting in a FACSAria to reduce debris/dead cells and enrich for specific immune populations prior to sequencing. Live cells were sorted into four populations: CD45^-^ (tumor and stroma), CD45^+^CD3^+^ (T cells), CD45^+^CD3^-^CD19^-^XCR1^+^ and CD45^+^CD3^-^CD19^-^CCR7^+^ (cDC1s and mregDCs), and CD45^+^CD3^-^CD19^-^XCR1^-^CCR7^-^ (other myeloid cells). These four populations were then recombined at a 1:2:1:1 ratio and submitted for sequencing (described below). Antibodies used for flow cytometry and FACS are listed in the Key resources table.

### Immunofluorescence

Tissue slides were thawed at RT in a sealed container for 5 min, transferred to acetone at −20°C for 5 min, and then left at RT again for 5-10 min to fully dry. Slides were then incubated in 0.05% Tween-20 in PBS (PBST) for 15 min, carefully dried with wipes, and tissue samples were delineated with oil marker. Thereafter, staining solutions were applied as a droplet (∼50-100μl) on top of each tissue sample to cover it completely and left in a humidified chamber during the incubation period to prevent it from drying out. Samples were blocked in 5% normal horse serum PBST for 30 min at RT. Labeled primary antibodies (Key resources table) were premixed in blocking solution according to each experimental setting. Blocking solution was discarded from the slides without washing, and then the antibody mix was added to the samples and incubated for 1 hour at RT. Slides were then washed twice in PBST in an orbital shaker. If a secondary antibody was needed, the slides were then incubated with the appropriate antibody mix for 1 hour at RT and washed twice in PBST. Slides were counterstained with DAPI for 5 min at RT and washed in PBS twice. Specimens were then preserved in Prolong© Gold Antifade mounting media. Tile-scanned images of the samples (9-12 tiles per image, 2-4 images per tumor condition) were obtained at 40X using the Zeiss Axio microscope for further analysis.

### Systemic secretion profiling

At the indicated timepoint, blood samples were collected by cardiac puncture under anesthesia as terminal surgery, and mice were subsequently euthanized for tumor processing. Samples were immediately transferred into tubes containing 10mL of 0.5M EDTA to prevent clotting and preserved on ice. Blood samples were then centrifuged at 1,500*g* for 10 min at 4°C. Plasma was collected, aliquoted as per instructions of Eve Technologies Corp., and frozen at −80°C until further processing. Samples were then submitted for multiplexed protein quantification using the Mouse Cytokine/Chemokine 44-plex Discovery Assay Array. For the assay, Eve Technologies calculated the requisite standard curves and returned quantified protein concentration measurements (pg/mL) for each sample. Out of range values were assigned the highest or lowest standard curve value per Eve Technologies instructions. Cytokines/chemokines on the panel which were either largely undetected or whose signal was overly saturated were removed for downstream analysis and visualization.

### Cellular indexing of transcriptomes and epitopes by sequencing (CITE-seq)

10,000 single cells from each condition were loaded onto the 10X Chromium System. Library preparation was conducted by the Yale Center for Genome Analysis (YCGA) using the Chromium Single Cell 5’ Reagent Kit with Feature Barcoding (10x Genomics) according to manufacturer instructions (see: [https://medicine.yale.edu/keck/ycga/sequencing/10x/singcellsequencing/]). Briefly, single cells were isolated in 1nL of Gel Beads-in-emulsion (GEMs) using the GemCode technology. Cell barcoding, lysis, and reverse transcription of mRNA occurred within each GEM. cDNA libraries were then generated using next-generation sequencing PCR amplification. cDNA quality control was performed with an Agilent TapeStation. Gene expression and cell surface protein libraries were generated for each sample. Libraries underwent paired-end sequencing on the NovaSeq 6000 platform (Illumina) by YCGA.

### Fabrication of microfluidic device for deterministic barcoding in tissue for spatial omics sequencing (DBiT-seq)

The comprehensive fabrication process, employing standard soft lithography, has been detailed in our previous publication.^77^ Briefly, high-resolution chrome photomasks with a customized pattern were printed and ordered from Front Range Photomasks (Lake Havasu City, AZ). Upon receipt, the masks underwent cleaning with acetone to remove any dirt or dust. Master wafers were then produced using SU-8 negative photoresist (SU-2010 or SU-2025) on silicon wafers following the manufacturer’s guidelines, with feature width of 50 µm, 20 µm, or 10 µm. The newly fabricated wafers were treated with chlorotrimethylsilane for 20 minutes to develop high-fidelity hydrophobic surfaces. Subsequently, polydimethylsiloxane (PDMS) microfluidic chips were fabricated through a replication molding process. The base and curing agents were mixed thoroughly with a 10:1 ratio following the manufacturer’s guidelines and poured over the master wafers. After degassing in the vacuum for 30 min, the PDMS was cured at 70°C for at least 2 hours. The solidified PDMS slab was cut out, and the inlets and outlets were punched for further use.

### DNA barcodes annealing

DNA oligos used in this study (Key resource table) were procured from Integrated DNA Technologies (IDT, Coralville, IA). (100 µM) and ligation linker (100 µM) were annealed at a 1:1 ratio in 2X annealing buffer (20 mM Tris-HCl pH 8.0, 100 mM NaCl, 2 mM EDTA) with the following PCR program: 95°C for 5 min, slow cooling to 20°C at a rate of −0.1°C/s, followed by 12°C for 3 min. The annealed barcodes can be stored at −20°C until use.

### Fixation and permeabilization, in situ polyadenylation, and reverse transcription

Tissue sections, obtained as detailed before, were retrieved from the −80°C freezer and equilibrated to RT for 10 min until all moisture dissipated. The tissue was permeabilized for 20 min at RT with 1% Triton X-100 in DPBS, followed by 0.5X DPBS-RI (1X DPBS diluted with nuclease-free water, 0.05 U/µL RNase Inhibitor) wash to halt permeabilization. The tissue slide was then air-dried and equipped with a PDMS reservoir covering the region of interest (ROI). *In situ* polyadenylation was performed using *E. coli* Poly(A) Polymerase. Initially, samples were equilibrated by adding 100 µL wash buffer (88 µL nuclease-free water, 10 µL 10X Poly(A) Reaction Buffer, 2 µL 40 U/µL RNase Inhibitor) and incubating at RT for 5 min. Following wash buffer removal, 60 µL of the Poly(A) enzymatic mix (38.4 µL nuclease-free water, 6 µL 10X Poly(A) Reaction Buffer, 6 µL 5U/µL Poly(A) Polymerase, 6 µL 10mM ATP, 2.4 µL 20 U/µL SUPERase•In RNase Inhibitor, 1.2 µL 40 U/µL RNase Inhibitor) was added to the reaction chamber and incubated in a humidified box at 37°C for 30 min. To remove excessive reagents, the slide was dipped in 50 mL DPBS and shake-washed for 5 min after the reaction. Subsequently, 60 µL of the reverse transcription mix (20 µL 25 µM RT Primer, 16.3 µL 0.5X DPBS-RI, 12 µL 5X RT Buffer, 6 µL 200U/µL Maxima H Minus Reverse Transcriptase, 4.5 µL 10mM dNTPs, 0.8 µL 20 U/µL SUPERase•In RNase Inhibitor, 0.4 µL 40 U/µL RNase Inhibitor) was loaded into the PDMS reservoir and sealed with parafilm. The sample was incubated at RT for 30 min and then at 42°C for 90 min, followed by a 50 mL DPBS wash as described before.

### Spatial barcoding with microfluidic devices

To ligate barcode A *in situ*, the first PDMS device was meticulously positioned atop the tissue slide, aligning the center channels over the ROI. The chip was imaged to record the positions for downstream alignment and analysis. Afterwards, an acrylic clamp was applied to firmly secure the PDMS to the slide, preventing any inter-channel leakage. The ligation mix, comprising 100 µL 1X NEBuffer 3.1, 61.3 µL nuclease-free water, 26 µL 10X T4 ligase buffer, 15 µL T4 DNA ligase, 5 µL 5% Triton X-100, 2 µL 40 U/µL RNase Inhibitor, and 0.7 µL 20 U/µL SUPERase•In RNase Inhibitor, was then prepared. For the barcoding reaction, 5 µL of the ligation solution, containing 4 µL ligation mix and 1 µL 25 µM DNA barcode A, was introduced into each of the 50 or 100 inlets. The solution was withdrawn to flow through the entire channel using a delicately adjusted vacuum. After a 30-minute incubation at 37°C, the PDMS chip was removed, and the slide was washed with 50 mL DPBS. Subsequently, the second PDMS device, featuring 50 or 100 channels perpendicular to the first PDMS, was attached to the ROI on the air-dried slide. A bright-field image was captured, and the ligation of barcode B set was performed similarly. Finally, after five flow-washes with 1 mL nuclease-free water to remove residual salt, the final scan was conducted to record the microchannel marks imprinted onto the tissue ROI.

### Tissue lysis and cDNA extraction

The barcoded tissue ROI was enclosed with a clean PDMS reservoir and securely clamped using acrylic chips. A 2X lysis buffer was prepared in advance, consisting of 20 mM Tris-HCl pH 8.0, 400 mM NaCl, 100 mM EDTA, and 4.4% SDS. For tissue digestion, 70 µL of the lysis mix (30 µL 1X DPBS, 30 µL 2X lysis buffer, 10 µL 20 µg/µL Proteinase K solution) was loaded into the PDMS reservoir, sealed with parafilm, and incubated in a humidified box at 55°C for 2 hours. After the reaction, the parafilm was removed, and all the liquid containing cDNA was collected into a 1.5mL DNA low-bind tube. Additionally, 40 µL of fresh lysis mix was loaded into the reservoir to collect any remaining cDNA material. The tissue lysate was incubated overnight at 55°C to completely reverse crosslinks, after which it could be stored at −80°C until the subsequent steps.

### cDNA purification, template switch, and PCR amplification

The cDNA was purified using 40 µL of Dynabeads MyOne Streptavidin C1 beads resuspended in 150 µL of 2X B&W buffer (10 mM Tris-HCl pH 7.5, 1 mM EDTA, 2 M NaCl). The mixture was incubated at RT for 60 min with rotation to ensure sufficient binding, followed by magnetic separation and two washes with 1X B&W buffer with 0.05% Tween-20, and an additional two washes with 10 mM Tris-HCl pH 7.5 containing 0.1% Tween-20. Streptavidin beads bound with cDNA molecules were then resuspended in 200 µL of TSO Mix (75 µL nuclease-free water, 40 µL 5X RT buffer, 40 µL 20% Ficoll PM-400, 20 µL 10mM dNTPs, 10 µL 200U/µL Maxima H Minus Reverse Transcriptase, 5 µL 40 U/µL RNase Inhibitor, 10 µL 100 µM TSO Primer). The template switch reaction was conducted at RT for 30 min and then at 42°C for 90 min with gentle rotation. After a single wash with 10 mM Tris-HCl pH 7.5 containing 0.1% Tween-20 and another wash with nuclease-free water, the beads were resuspended in 200 µL of PCR Mix (100 µL 2X KAPA HiFi HotStart ReadyMix, 84 µL nuclease-free water, 8 µL 10 µM PCR Primer 1, 8 µL 10 µM PCR Primer 2). This suspension was then distributed into PCR stripe tubes. An initial amplification was conducted with the following PCR program: 95°C for 3 min, cycling five times at 98°C for 20 s, 63°C for 45 s, 72°C for 3 min, followed by an extension at 72°C for 3 min and 4°C hold. Following magnetic removal of the beads, 19 µL of the PCR solution was combined with 1 µL 20X EvaGreen for quantitative real-time PCR (qPCR) analysis using the same program. The remaining samples underwent further amplification, with the cycle numbers determined by 1/2 of the saturated signal observed in qPCR results. The PCR product was then purified using SPRIselect beads at a 0.8X ratio, adhering to the standard manufacturer’s instructions. The resulting cDNA amplicon underwent analysis using a TapeStation system with D5000 DNA ScreenTape and reagents. This stage provides a secure stopping point, allowing the sample to be stored at −20°C until the next steps.

### rRNA removal, library preparation, and sequencing

The SEQuoia RiboDepletion Kit was employed to eliminate fragments derived from rRNA and mitochondrial rRNA from the amplified cDNA product, following the manufacturer’s guidelines. Based on the TapeStation readout profile, 20 ng of cDNA was used as the input amount, and three rounds of depletion were performed. Subsequently, 7 cycles of the aforementioned PCR program were executed to directly ligate sequencing primers, using a 100 µL system consisting of 50 µL 2X KAPA HiFi HotStart ReadyMix, ∼42 µL solution from the rRNA removal step, 4 µL 10 µM P5 Primer, and 4 µL 10 µM P7 Primer. The resulting library underwent purification using SPRIselect beads at a 0.8X ratio, quality control checked using TapeStation, and was then sequenced on an Illumina NovaSeq 6000 Sequencing System with a paired-end 150bp read length.

## QUANTIFICATION AND STATISTICAL ANALYSIS

### CITE-seq preprocessing and annotation

Gene expression reads were aligned to the reference genome mm10-2020-A using the cellranger multi pipeline from the CellRanger software v6.1.0 (10x Genomics) to generate cell-by-gene matrices. Antibody capture reads were aligned to a reference csv summarizing the set of Feature Barcode reagents used in the experiment, also using the cellranger multi pipeline from the CellRanger software v6.1.0 (10x Genomics) to generate cell-by-antibody matrices. Downstream analyses were performed using the Scanpy package in Python.^71^ Single cells were filtered to remove doublets, dead cells, and otherwise low-quality cells. Cells with total transcript counts less than 1,000 or greater than 60,000-70,000, with total detected genes less than 200 or greater than 6,500-8,000, or with mitochondrial gene counts exceeding 15% were removed from the dataset for downstream analyses. As cells from different tumors were labeled with different hashtag antibodies prior to cell pooling and single cell isolation, correctly captured singlets could be identified by their expression of only one hashtag antibody. Doublets and multiplets were then easily identified by their expression of two or more hashtag antibodies.^78^ The log_10_-transformed expression profiles for each hashtag antibody were fit with Gaussian mixture models to establish detection thresholds. Single cells then found to express more than one hashtag antibody were filtered from the dataset for downstream analysis. Dimensionality reduction and visualization were completed with the Scanpy implementation of Uniform Manifold Projection and Approximation (UMAP).^79^ To complete cell type annotation, we adapted a previously published pipeline^80^ to use a simple feedforward neural network (NN), as our group has done previously.^20,81,82^

### Cell-cell communication inference

To infer cell-cell communication from scRNA-seq, we used the Niche Interactions and Communication Heterogeneity in Extracellular Signaling (NICHES) method.^10^ While most methods to construct scRNA-seq-derived cell-cell communication network require a user’s data to be pre-clustered,^4^ NICHES infers ligand-receptor communication at the level of pairs of single cells sampled from the dataset, revealing previously masked heterogeneity in predicted cell -cell signaling. To combat the propagation of dropout events, we first imputed the scRNA-seq data using the adaptively thresholded low rank approximation (ALRA) method.^83^ We then used NICHES with default parameters to construct matrices of ordered cell-cell pairs by known ligand-receptor axes, where matrix entries were the product of ligand and receptor expression in the corresponding pair of single cells. Cell-cell pairs were limited to disallow pairing between cells from different replicate tumors, and literature-supported ligand-receptor pairs were sourced from the OmniPath database.^74^ We further limited our analyses to consider only cell-cell communication between macrophages, DC subsets, and T cell subsets because these cell populations were directly downstream of therapy. ALRA and NICHES are both available as open-source software packages in R.

### Differential abundance testing

We used the Milo method^41^ (as implemented in Python) for differential abundance testing. Milo outperforms competing methods^62,63^ in flexibility of experimental design and in accuracy and robustness despite technical variability. Hashtag antibody expression was used to distinguish replicates within each experimental condition, and differentially abundant neighborhoods (spatialFDR < 0.1) were identified with default parameters. For ICB only-treated samples, differential abundance (DA) testing was performed w.r.t. percentage of maximum treatment dose (continuous covariate where control = 0, ICB lo = 25, ICB hi = 100). For NICHES outputs comparing control, ICB lo-, and ICB lo + CD40ag-treated samples, DA testing was performed w.r.t. CD40ag inclusion (discrete covariate where control = 0, ICB lo = 0, CD40ag = 1, ICB lo+CD40ag = 1). For NICHES outputs comparing ICB lo + CD40ag treatment at days 8 and 10, DA testing was performed w.r.t. days post-therapy (continuous covariate where day 8 = 1, day 10 = 3). Differentially abundant neighborhoods were then clustered using the Louvain algorithm as was done in Dann et al.^41^, with adjacent neighborhoods defined as those with at least two overlapping single cells or cell-cell pairs and an absolute difference in log-fold change below a specified threshold depending on the comparison. Extracted neighborhood clusters were analyzed using custom scripts in Python. Differentially expressed genes (DEGs) between neighborhood clusters were identified using the rank_genes_groups function from the Scanpy module in Python with default parameters (log_2_FC > 1.5 and adjusted p-value < 0.05).

### Fluorescent image segmentation, annotation, and quantification

Tile scan images, obtained as detailed previously, were stitched using the Zen Pro v2.3 software package, and uploaded onto QuPath^84^ (v0.5.0) for segmentation and classification. Cells were segmented using the DAPI nuclear staining and then classified by cell type based on thresholded fluorescence intensity of each channel. The XY coordinates corresponding to cells of each cell type were then used for Euclidean distance calculations in MATLAB.

### DBiT-seq preprocessing and cell type scoring

Downstream analyses were performed using Seurat v4.3.0^72^ and custom scripts in R. Data were normalized using the SCTransform function. Genes detected in fewer than 2 spatial spots per sample were filtered out. To calculate cell type enrichment scores for each spot, we first identified the top 20 DEGs for each cell type captured in the scRNA-seq data (log_2_FC > 1.5 and adjusted p-value < 0.05; see “CITE-seq analysis”) and which were consistently detected across DBiT-seq samples. Cell type signature scoring was then accomplished using the AddModuleScore function (Seurat) using 100 randomly selected control genes. Macrophage signature genes included *Arhgef37, Msr1, Lyz2, Cybb, C1qb, Vcan, F13a1, Maf, C1qc, Ms4a6c, Lpl, Cmklr1, Ctsc, C1qa, Ccl2, Ccr2, Fn1, Lrp1, Arg1,* and *Ms4a8a*. mregDC signature genes included *Tmem150c, Stap2, Slco5a1. Dscaml1. Chst3, Dpysl5, Oprd1, Fscn1, Ccr7, Cacnb3, Cd200, Mreg, Socs2, Synpo2, Ramp3, Il4i1, Adcy6, Sema7a, Casc1,* and *Efna2*. Treg signature genes included *Lrrc32, Itgae, Ikzf2, Itgb8, Neb, Ecm1, Lgmn, Ankrd55, Maf, Ass1, Gpld1, Tnfrsf9, Cd5, Lamc1, Ncmap, Nid2, Capg, Glrx, Arl5a,* and *Dusp4*. CD8+ T cell signature genes included *Tmem108, Jaml, Cd3d, Gramd3, Nsg2, Fam241a, Hdgfl3, Actn1, Tcf7, Ccr7, Sidt1, Ifit3, Pde7a, Ms4a6b, Bcl2, Rnf213, Dnajc15, Epsti1, Slco3a1,* and *Evl*.

### Statistical analysis

Data were presented as mean ± SD unless otherwise specified. Statistical analysis was generally performed by Wilcoxon rank-sum testing, or as specified in the figure legends. Values were considered significant at *P* < 0.05. All analyses were performed using either custom python scripts or GraphPad Prism (Version 10.0.3).

## SUPPLEMENTAL INFORMATION

Document S1. Figures S1-S10 and Table S1.

Supplemental File 1. Excel file containing full suite of differential expression results for differentially abundant neighborhood clusters of cell-cell communication, related to Fig. 3.

Supplemental File 2. Excel file containing full suite of differential expression results for differentially abundant neighborhood clusters of cell-cell communication, related to Fig. 4.

Supplemental File 3. Excel file containing full suite of differential expression results for differentially abundant neighborhood clusters of cell-cell communication, related to Fig. 5.

Supplemental File 4. Excel file containing full suite of differential expression results for ‘durable’ neighborhood clusters of cell-cell communication, related to Fig. 7.

