## Supplementary material for "Mapping intratumoral myeloid-T cell interactomes at single-cell resolution reveals targets for overcoming checkpoint inhibitor resistance": DocumentS1

*Figure S1: Treatment-induced changes in size, cell type composition, and phenotype in 8 day YR1.7 tumors*

A

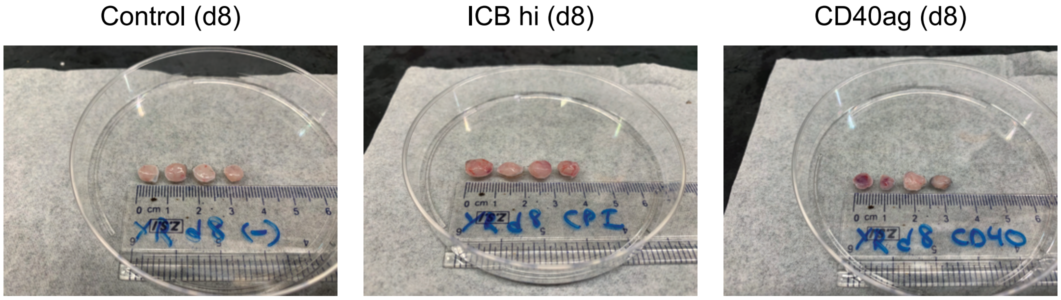


B

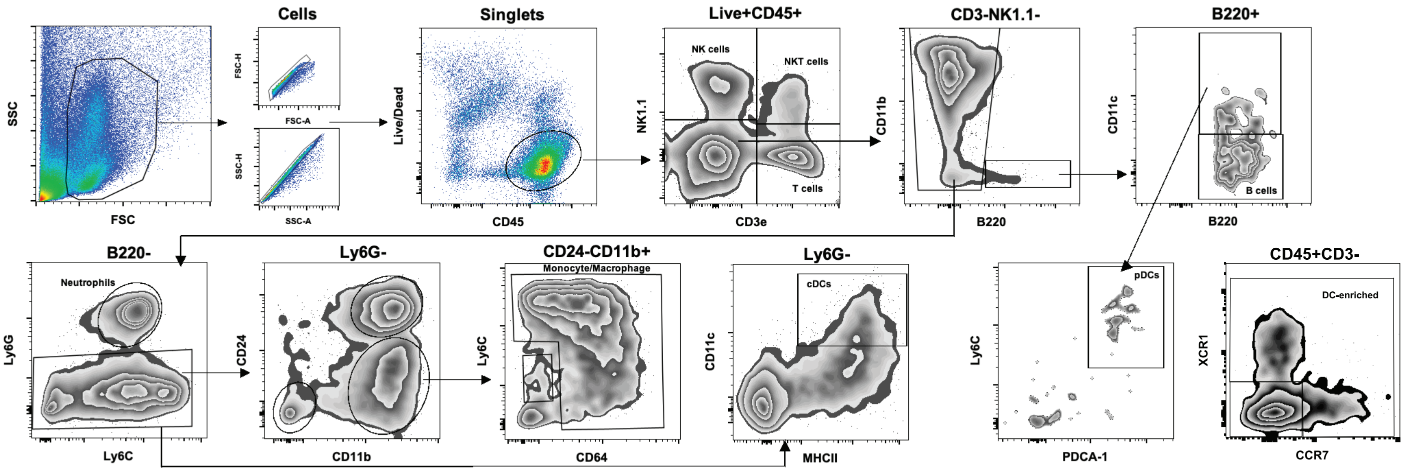


C

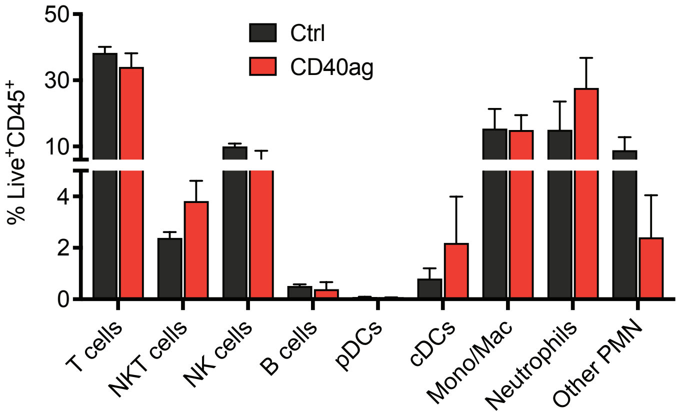


Related to Fig. 1. (A) Images of individual 8 day YR1.7 tumors either left untreated (left), or treated with ICB hi (center) or CD40ag monotherapy (right) 24 hours prior to excision.
(B) Flow cytometry gating strategy for analyzing immune populations in YUMMER1.7 tumors.
(C) Bar graphs describing quantification of immune populations as sorted in (B) from control (black) and CD40ag-treated (red) tumors as a percentage of total Live$^{+}$CD45$^{+}$ cells. Data is represented as mean ± SEM.

*Figure S2: Gene expression patterns were consistent across replicate tumors, confirmed cell types immediately downstream of therapy*

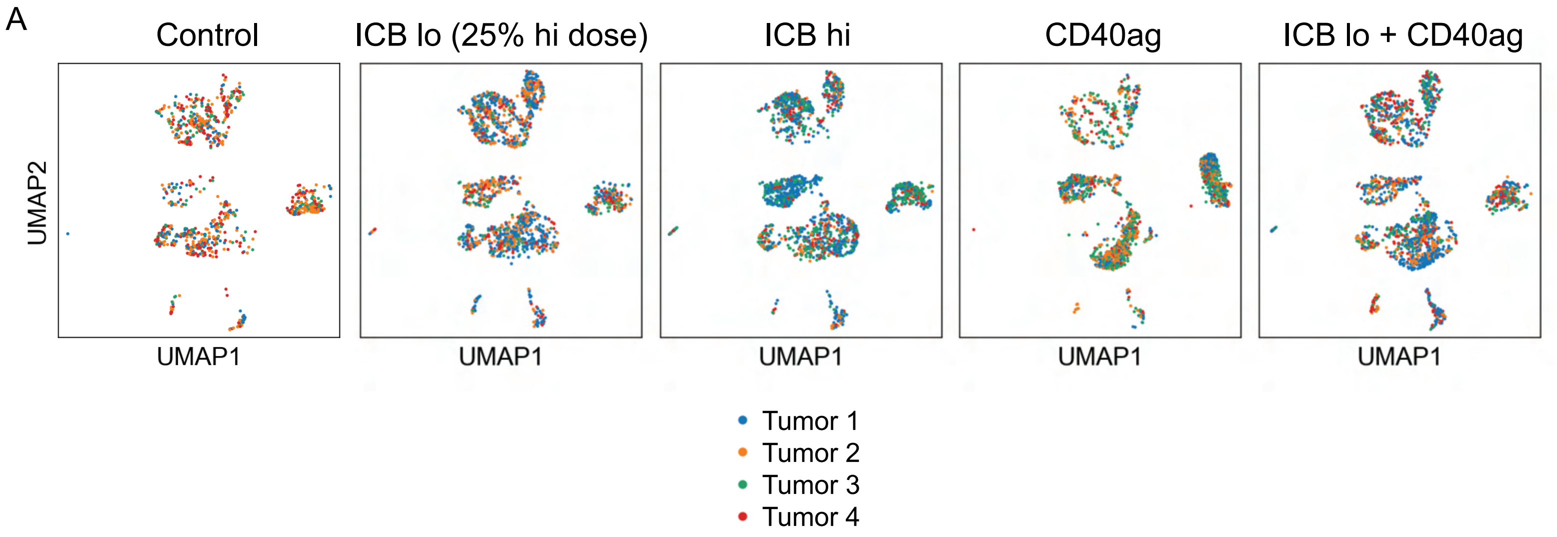


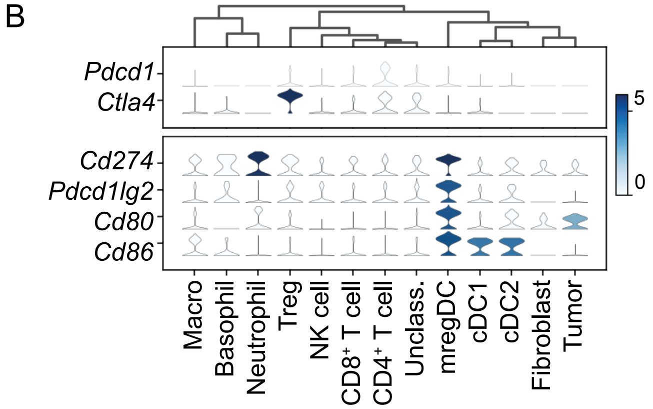


Related to Fig. 2. (A) UMAP embedding of control, ICB lo-treated, ICB hi-treated, CD40ag-treated, and ICB lo + CD40ag-treated scRNA-seq samples separated by treatment condition and colored by individual tumor (4 tumors per group). Single cells from identically-treated tumors were generally well-mixed in two-dimensional space.
(B) Expression of genes which encode for ICB targets (top) and their cognate ligands (bottom) across scRNA-seq samples by cell type.

*Figure S3: NICHES and Milo uncover similar axes of acute myeloid-T cell crosstalk across tumor models treated with CD40ag monotherapy*

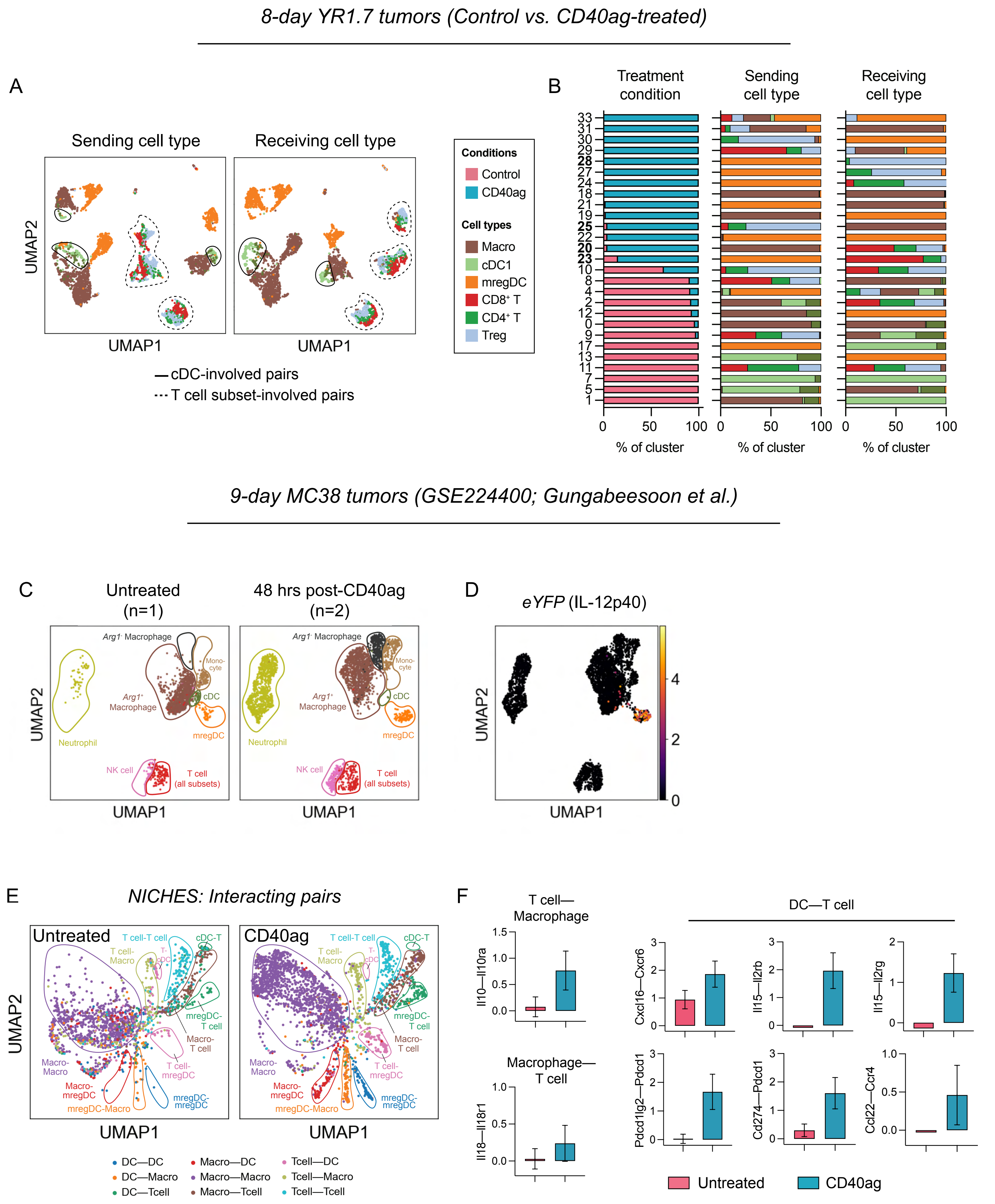

Related to Fig. 3. (A) Integrated UMAP embedding of NICHES interactomes in Fig. 3A colored by sending (left) and receiving (right) cell subset.

(B) Stacked bar graph summarizing the cell-cell pairs which make up each differentially abundant cluster in Fig. 3C by (left) treatment condition, (center) sending cell type, and (right) receiving cell type.

(C) UMAP embedding of publicly available scRNA-seq data (GSE224400) of untreated (left) and CD40ag-treated (right) MC38 tumors 9 days post-tumor initiation and 48 hours post-treatment where applicable. Data is colored by cell type annotations from the original publication.

(D) UMAP embedding as in (C) colored by expression of eYFP (IL-12p40).

(E) UMAP embedding of NICHES interactomes for control (n=1 tumor) and CD40ag monotherapy-treated (n=2 tumors) MC38 samples colored by broad cell type pair. Tumors were collected 9 days post-inoculation and 48 hours post-treatment where applicable.

(G) Bar graphs comparing predicted interaction scores across untreated (pink) and CD40ag monotherapy-treated (blue) T cell-macrophage (top left), macrophage-T cell (bottom left), and DC-T cell (right) pairs for selected ligand-receptor axes. Bar graphs are represented as mean interaction score ± 95% CI (calculated with GraphPad).

*Figure S4: ICB rewires myeloid-T cell signaling and cell subset abundance within 24 hours of treatment*

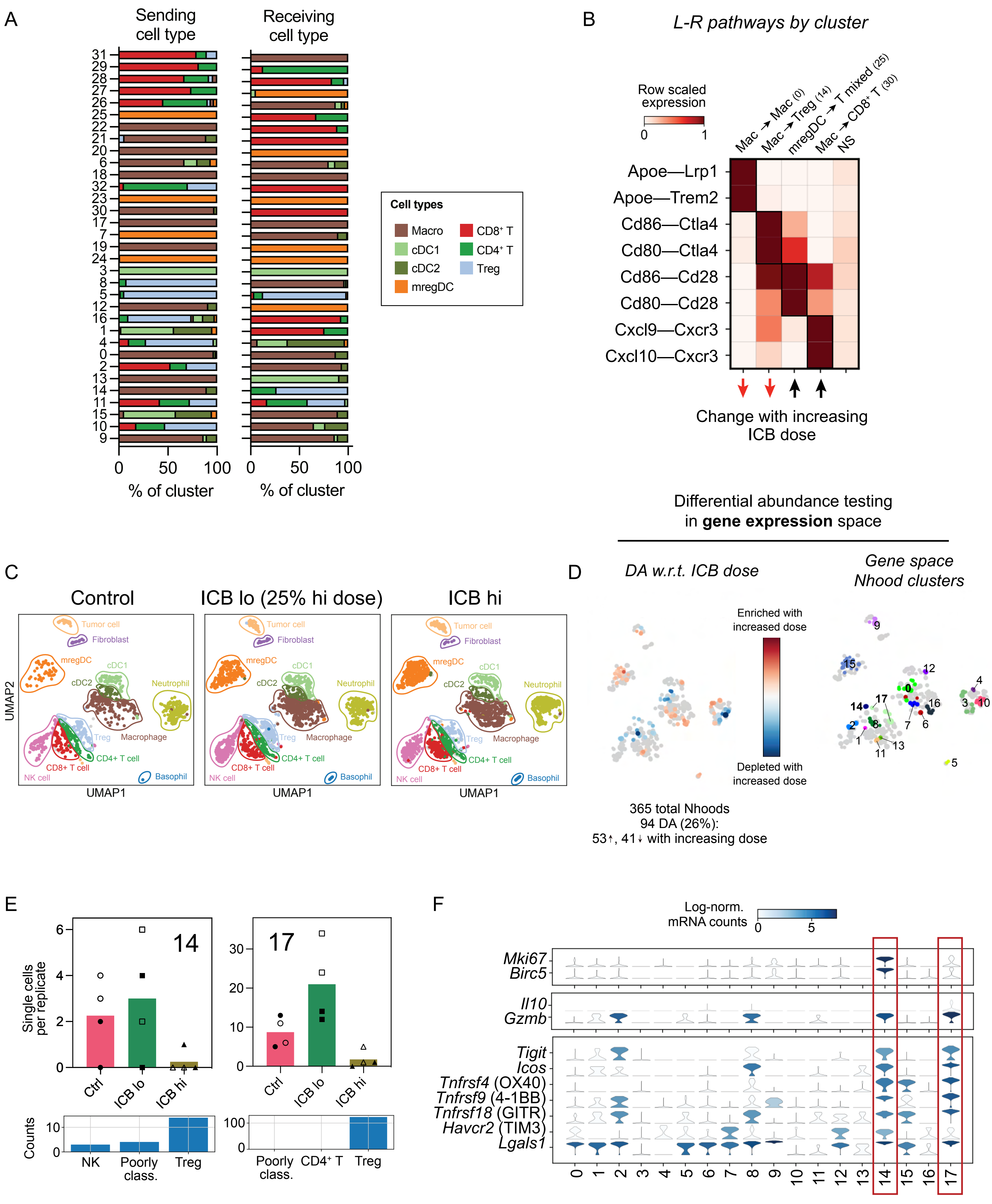

Related to Fig. 4. (A) Stacked bar graphs summarizing the cell-cell pairs which make up each differentially abundant cluster in Fig. 4C by sending cell type (left) and receiving cell type (right).
(B) Heatmap of per-row scaled interaction scores for selected differentially predicted ligand-receptor axes (y-axis) across clusters 0, 14, 25, and 30 and compared to all non-DA neighborhoods (i.e., NS) (x-axis).
(C) UMAP embedding of control (left), ICB lo-treated (center), and ICB hi-treated (right) samples (n=4 tumors per group) separated by treatment condition and colored by cell type as assigned by a neural network-base classifier.
(D) UMAP embedding of (left) 365 Milo-assigned ‘neighborhoods’ (nhoods) of single cells colored by differential abundance result w.r.t. ICB dose, and of (right) DA nhoods (n=94) colored by grouping for downstream analyses. Grey nhoods were not DA (n=271), while blue (n=41) and red n=53) nhoods decreased and increased in frequency, respectively, with ICB dose.
(E) Bar graphs summarizing number of single cells averaged across replicates (y-axis) per treatment (x-axis) (top) and distribution of annotated cell types (bottom) for DA clusters 14 and 17. Markers distinguish tumor samples from different mice (n=2 per experimental group).
(F) Expression of proliferation markers *Mki67* and *Birc5* (top), immunosuppressive signal-encoding genes *Il10* and *Gzmb* (middle), and immune checkpoints *Tigit, Icos, Tnfrsf4, Tnfrsf9, Tnfrsf18, Havcr2,* and *Lgals1*. Violin shape represents the distribution of gene expression across single cells in each cluster, while the color fill represents average gene expression of each gene across each cluster.

*Figure S5: Immunotherapy differentially alters immune composition and relative spatial orientation in the TME*

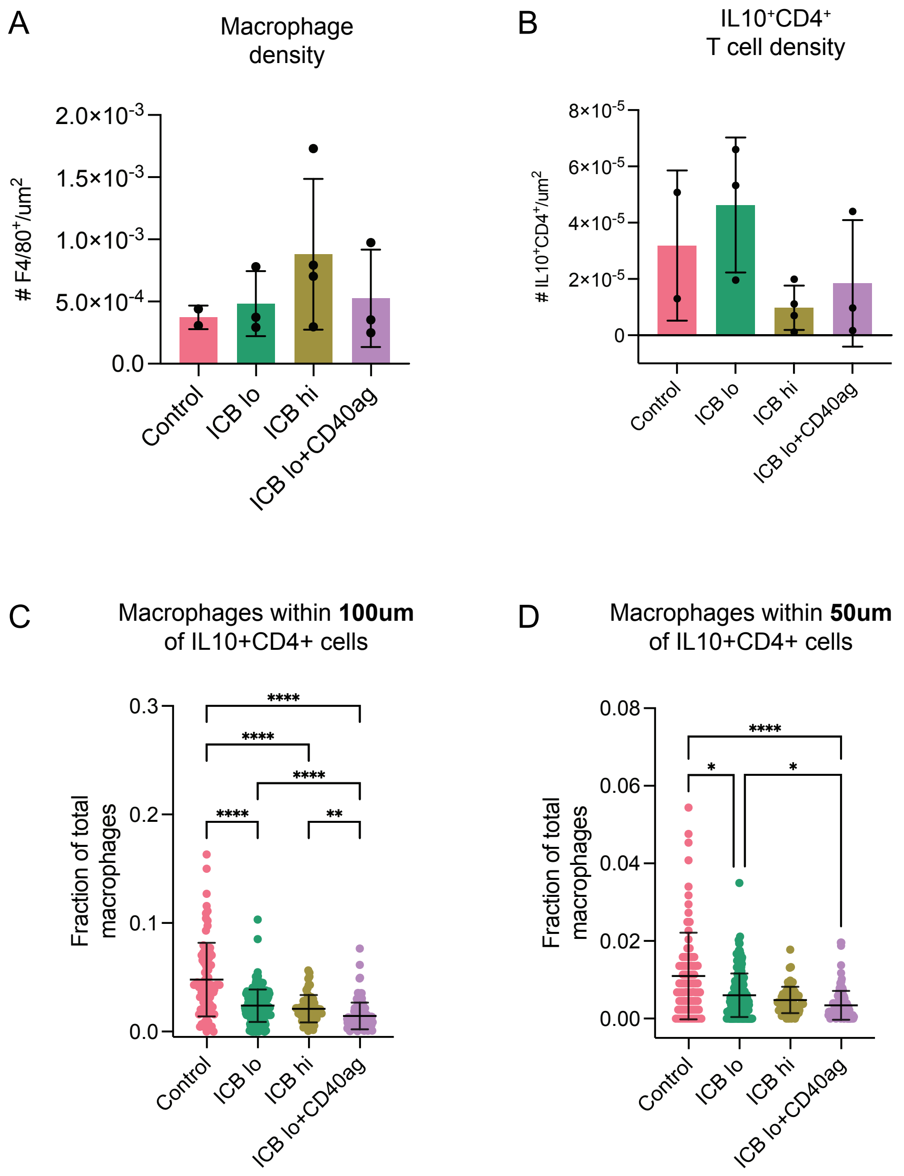


Related to Fig. 4. (A-B) Bar graphs demonstrating quantification of (A) CD4$^{+}$IL-10$^{+}$ T cell and (B) macrophage (F4/80$^{+}$) density (absolute number of cells divided by tumor area [um$^{2}$])from control (pink), ICB lo- (green), ICB hi- (yellow), and ICB lo + CD40ag-treated (purple) tumor slices (n=2-4 regions from 1 tumor per treatment condition). Summary data is presented as mean ± SD.

(C-D) Scatter plots demonstrating the fraction of total macrophages (F4/80$^{+}$) within (C) 100 um and (D) 50 um of CD4+IL-10+ T cells from control (pink), ICB lo- (green), ICB hi- (yellow), and ICB lo + CD40ag-treated (purple) tumor slices. Summary data is presented as mean ± SD. ****$p < 0.0001$ by Kruskal-Wallis test (calculated in GraphPad).

*Figure S6: NICHES + Milo identify axes of cell-cell communication specific to combinatorial therapy*

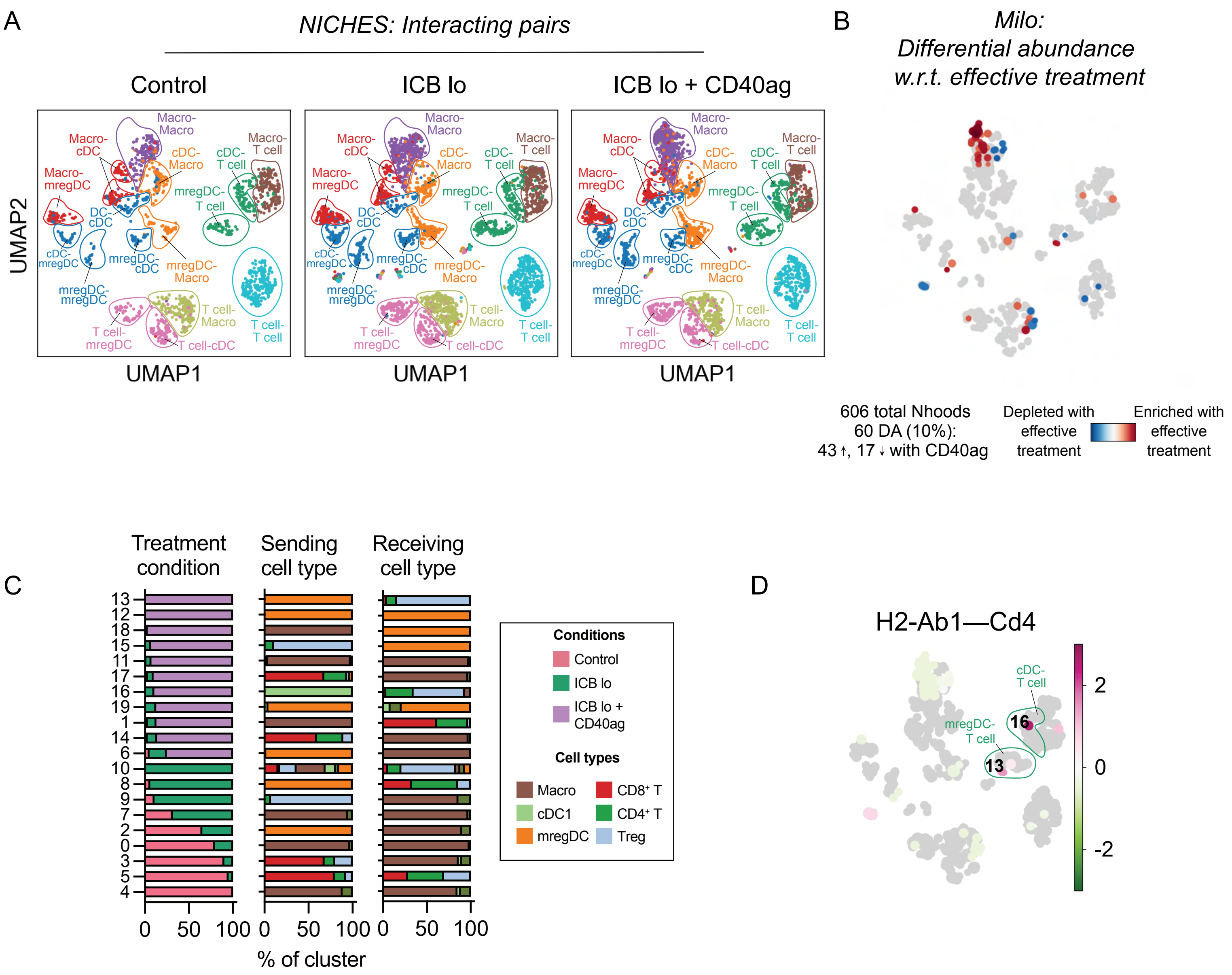


Related to Fig. 5. (A) UMAP embeddings of NICHES interactomes from control, ICB lo- and ICB lo + CD40ag-treated samples separated by treatment condition and colored by broad cell type pair; and (B) UMAP embedding of 606 Milo-assigned 'neighborhoods' (nhoods) of cell-cell pairs colored by differential abundance result w.r.t. effective treatment [i.e., Control, ICB lo = ineffective (0); ICB lo + CD40ag = effective (1)]. Grey nhoods were not differentially abundant (n=546), while blue (n=17) and red (n=43) nhoods decreased and increased in frequency, respectively, with effective treatment. (C) Stacked bar graphs summarizing the cell-cell pairs which make up each differentially abundant cluster in Fig. 5A by treatment condition (left), sending cell type (center), and receiving cell type (right). (D) UMAP embedding as in (A) colored by predicted interaction score along H2-Ab1 – Cd4 as averaged across single cell-cell pairs in each nhood.

*Figure S7: CD40ag-induced IL-12 signaling impacts acute trends in tumor growth but not long-term survival*

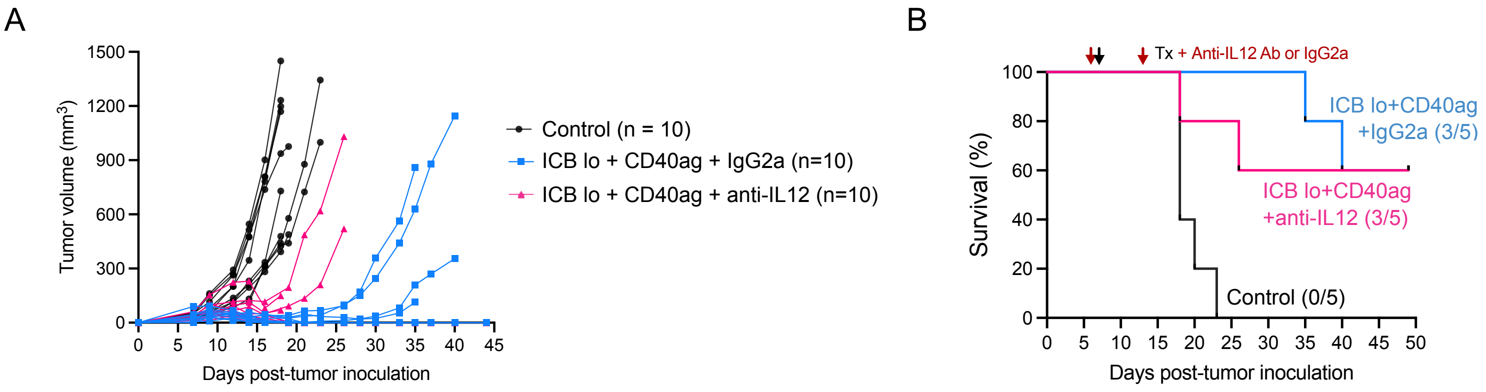


Related to Fig. 5. (A) Tumor volume ($mm^3$) curves over time for YUMMER1.7-bearing mice (2 tumors per mouse) either left untreated (n=10), treated with ICB lo + CD40ag + IgGa (n=10), or treated with ICB lo + CD40ag + IL-12 blockade (n=10).

(B) Corresponding Kaplan-Meier survival curves for tumor volume data shown in (A).

*Figure S8: CD40ag in combination with ICB primarily alters myeloid heterogeneity*

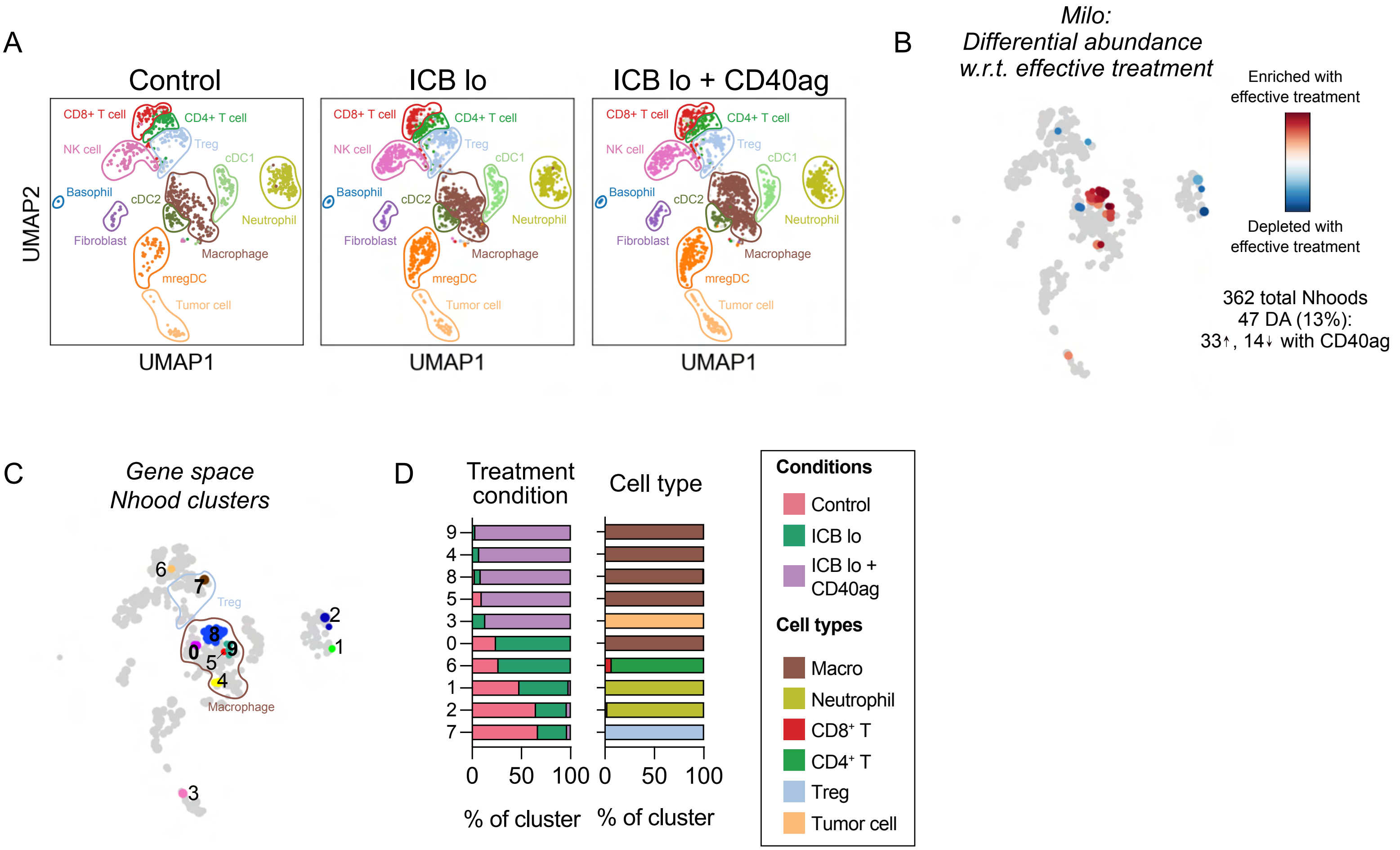


Related to Fig. 5. (A) UMAP embedding of control (left), ICB lo-treated (center), and ICB lo+CD40ag-treated (right) samples (n=4 tumors per group) separated by treatment condition and colored by cell type as assigned by a neural network-based classifier.

(B) UMAP embedding as in (A) of 362 Milo-assigned 'neighborhoods' (nhoods) of single cells colored by differential abundance result w.r.t. effective treatment [i.e., Control, ICB lo = ineffective (0); ICB lo + CD40ag = effective (1)]. . Grey nhoods were not DA (n=315), while blue (n=14) and red (n=33) nhoods decreased and increased in frequency, respectively, with effective treatment.

(C) UMAP embedding of as in (B) of DA nhoods (n=47) colored by grouping for downstream analyses.

(D) Stacked bar graphs summarizing the cells which make up each differentially abundant cluster in (C) by treatment condition (left) and cell type (right).

*Figure S9: Blocking IL-12/IL-18 signaling alters peripheral blood secretion activated by combinatorial therapy*

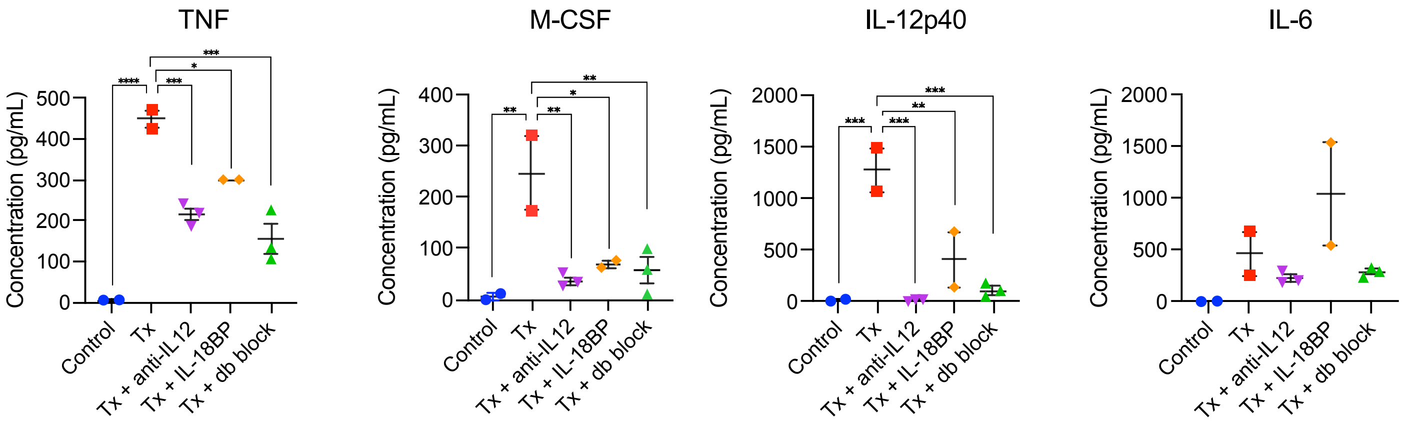


Related to Fig. 5. Example plots of peripheral blood cytokine expression levels in YUMMER1.7-bearing mice for TNF, M-CSF, IL-12p40, and IL-6. Mice were either left untreated (blue) or were treated with ICB lo + CD40ag (abbrev. Tx, red), Tx + anti-IL12 (purple), Tx + IL-18BP (orange), or Tx + anti-IL12 + IL-18BP ('db block', green). Summary data is presented as mean ± SEM. *p < 0.05, **p < 0.01, ***p < 0.001, ***p < 0.0001 by ordinary one-way ANOVA with Dunnett's multiple comparisons testing (calculated in GraphPad). Only showing significant results for comparisons between Tx and other conditions.

*Fig. S10: Macrophages and DCs do not colocalize similarly with T cell subsets 24 hours post-therapy*

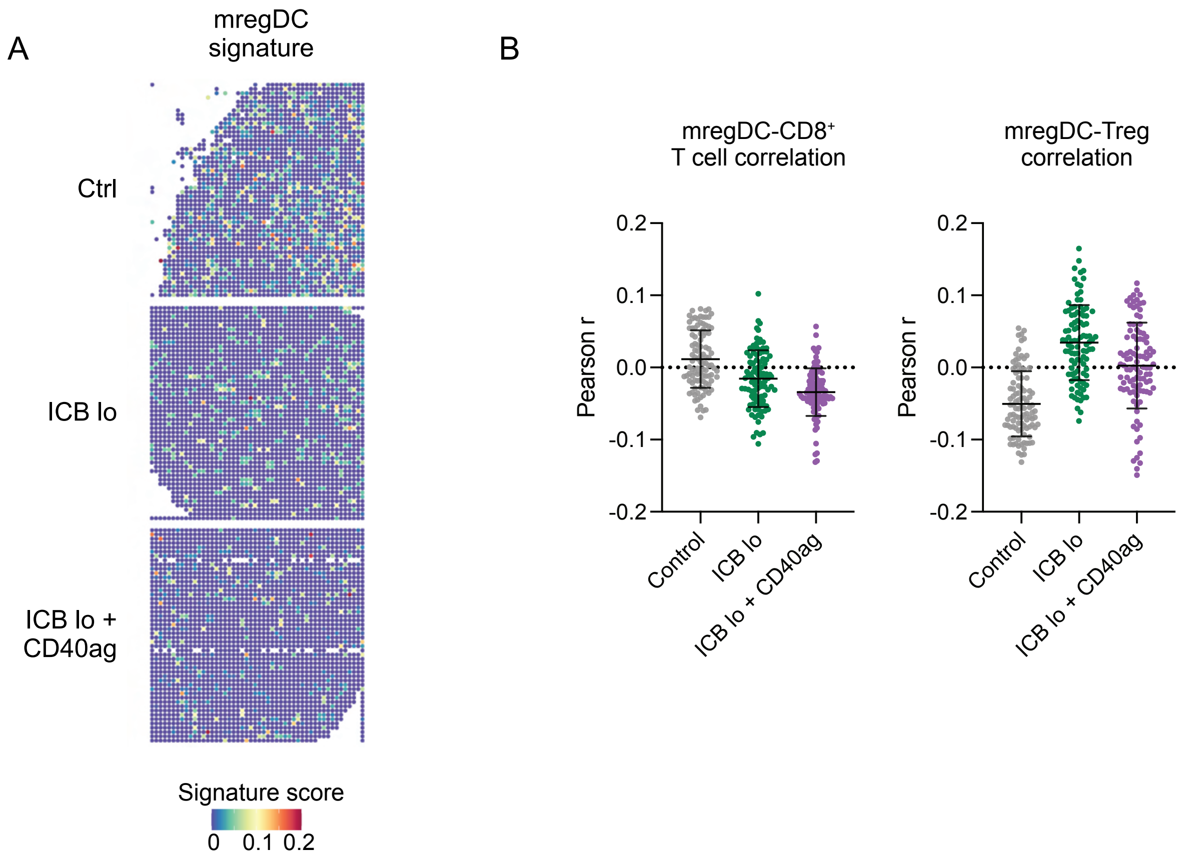


Related to Fig. 6. (A) mregDC signature scores for each pixel in the spatial transcriptomics spot map. (B) Scatter plots demonstrating per-pixel Pearson product-moment correlation coefficients between (left) mregDC and $CD8^+$ T cell signature scores and (right) mregDC and Treg signature scores for 100 20x20 grids randomly sampled from each DBiT-seq sample (control – grey, ICB lo – green, ICB lo + CD40ag – purple). Summary data is presented as mean ± SD.

Table S1. Summary of all differentially abundant neighborhood clusters in Fig. 5A.

| Cluster no. | Enriched for: | Sending cell type | Receiving cell type | Key signaling axes | Hypothesized function | Supporting references |
|---|---|---|---|---|---|---|
| 0 | Untreated | Macrophage | Macrophage | Apoe—Trem2<br>Spp1—Cd44 | Anti-inflammatory macrophage activation, tumor growth, anti-PD-1 resistance | 1,2 |
| 1 | Tx | Macrophage | $CD8^{+}$/<br>$CD4^{+}$ T | Described in Fig. 5 | | |
| 2 | Untreated | mregDC | Macrophage | Ccl5—Ccr(1,5)<br>Cx3cl1—Cx3cr1 | Chemotaxis, myeloid infiltration into the TME | 3 |
| 3 | Untreated | Mixed T cell | Macrophage | Ifng—Ifngr(1,2)<br>Adam10—Trem2<br>Cd47—Sirpa | Dual pro- and anti-inflammatory macrophage polarization | 4 |
| 4 | Untreated | Macrophage | Macrophage | Ccl12—Ccr2<br>Ccl7—Ccr(1, 2,5) | Monocyte/macrophage recruitment into the tumor | 5,6 |
| 5 | Untreated | $CD8^{+}$ T | Mixed T cell | Ifng—Ifngr1<br>Cd274—Pdcd1 | Suppression of T cell proliferation and IFNg production, tumor tolerance | 7 |
| 6 | Tx | mregDC | Macrophage | Ccl5—Ccr(1,5)<br>Anxa2—Tlr2 | Chemotaxis, proinflammatory macrophage polarization, proinflammatory interleukin secretion | 8 |
| 7 | ICB lo | Macrophage | Macrophage | Apoe—Trem2<br>C3—C3ar1 | Anti-inflammatory macrophage activation, regulation of TIM-3 expression | 1,9 |
| 8 | ICB lo | mregDC | Mixed T cell | Cxcl16—Cxcr6<br>Cd80/86—Cd28<br>Il15—Il2r(b,g) | Positioning and survival of CTLs, sustained tumor control | 10 |

| | | | | | | |
|---|---|---|---|---|---|---|
| 9 | ICB lo | Treg | Macrophage | Described in Fig. 5 | | |
| 10 | ICB lo | Mixed | Mixed | - | - | - |
| 11 | Tx | Macrophage | Macrophage | Ccl2—Ccr2<br>Ccl(2,4, 12)—Ccr(1,5) | Chemotaxis, myeloid infiltration into the TME, macrophage maturation | [11] |
| 12 | Tx | mregDC | mregDC | Sema7a—Itgb1 | DC migration | [12] |
| 13 | Tx | mregDC | Treg | Described in Fig. 5 | | |
| 14 | Tx | $CD8^+/CD4^+$ T | Macrophage | | | |
| 15 | Tx | Treg | mregDC | | | |
| 16 | Tx | cDC1 | Mixed T cell | Described in Fig. S6 | | |
| 17 | Tx | $CD8^+/CD4^+$ T | Macrophage | Ccl5—Ccr(1,5)<br>Adam10—Trem2<br>Cd47—Sirpa | T cell survival, modulation of phagocytosis | [13,14] |
| 18 | Tx | Macrophage | mregDC | Il18—Cd48<br>Fn1—Cd44 | Chemotaxis, cell-cell adhesion | [15–17] |
| 19 | Tx | mregDC | mregDC | Sema7a—Itgb1 | DC migration | [12] |
